# Confidence: Its Measurement, Relation to Behavior, and Neural Representation

**DOI:** 10.64898/2026.11.05.764725

**Authors:** Mizuki Tsutsumi, Takeshi Ogawa, Tomohiko Kishi, Motoaki Kawanabe, Toshikazu Kuroda

**Affiliations:** Department of Dynamic Brain Imaging, Cognitive Mechanisms Laboratories, Advanced Telecommunications Research Institute International, 2-2-2, Hikaridai, Seika-cho, Soraku-gun, Kyoto 619-0288, Japan

## Abstract

Confidence (certainty/uncertainty about one’s own decision) has attracted considerable attention in psychology, neuroscience, and artificial intelligence. Yet its relation to behavior remains unclear: Some treat confidence as a cause of behavior whereas others view it as a state controlled by environmental variables. The unobservable nature of confidence makes its measurement validation and analyses difficult. To tackle these issues, we developed a novel procedure combining a decision-making task, risk choices, and a post-task open-ended questionnaire. Point-gain/loss amounts for correct/incorrect decisions varied depending on choices between high-risk/high-return and low-risk/low-return. In the questionnaire, 90% of 16 participants reported feelings interpretable as certainty/uncertainty when making those risk choices despite no instruction about confidence, whereas 8% did so when the choices led to identical outcomes/returns. This suggests confidence is a state controlled by environmental contingencies rather than a cause of behavior. We also decoded changes in the state from electroencephalography and functional magnetic resonance imaging.

---

Confidence refers to a feeling of certainty/uncertainty about other aspects of one’s own cognition such as perception, memory, and decision. As part of metacognition, this concept has attracted considerable attention across psychology, neuroscience, and artificial intelligence^1,2,3,4,5^. Despite extensive research, however, the relation between confidence and behavior remains unclear, especially with regard to whether one causes the other or the two just covary in response to common environmental variables.

A prominent view treats confidence as a *cause* of behavior. According to this perspective, confidence guides exploration, information seeking, learning, and decision strategies^6,7,8,9^. An alternative view is that confidence is not a cause of behavior but rather a description of an organism’s *state* controlled by environmental contingencies and task conditions^10,11,12,13^. From this perspective, the environment is a cause of both confidence and behavior, with the latter two just covarying. This distinction parallels the classic issue critically discussed in a seminal paper in psychology by MacCorquodale and Meehl (1948) ^14^: whether a particular concept is a *hypothetical construct* (i.e., an inferred entity/process which can causally affect behavior beyond effects of observable events) or an *intervening variable* (i.e., a summary description of relations among observable events without independent causal status). Thus, a fundamental question remains unresolved: Is confidence a cause of behavior or just a state generated by environmental variables?

Progress on this question has been hindered by the fact that confidence is not directly observable by a third party. Existing studies typically infer confidence either from self-ratings of confidence or from behavioral indices such as wagering and uncertainty responses^4,15,16^. Although these approaches have proven useful, they face a common challenge: measurement validity. In particular, it remains difficult to determine whether confidence measured those ways reflects what participants actually feel and, if so, how the feeling is related to behavior.

In the present study, we developed a novel procedure designed to tackle these issues directly. Participants completed a decision-making task, which involved risk choices (high-risk/high-return versus low-risk/low-return), and subsequently provided open-ended descriptions of what they felt when making those risk choices. Importantly, we gave them no instruction about confidence throughout the experiment. Moreover, the same participants also completed a control condition in which the two risk choices led to identical outcomes/returns (i.e., the contingency between risk choices and their differential outcomes was removed) with all other aspects of the task held constant. If references to certainty/uncertainty appear in the presence of the contingency and disappear in its absence, this would suggest that confidence is a state generated by environmental contingencies rather than a cause of behavior.

During the above decision-making task, we also conducted a simultaneous recording of electroencephalography (EEG) and functional magnetic resonance imaging (fMRI). Previous studies have identified several neural activities associated with confidence judgements, including centroparietal accumulation-like activity^17^, the error-related negativity/positivity in EEG^18,19,20^ and the dorsolateral and ventromedial prefrontal cortices and parietal regions in fMRI ^21,22,23,24^. Here, instead of treating neural correlates as evidence for confidence (i.e., a common logical fallacy in neuroimaging^25^), we asked how confidence, which was independently validated through the present decision-making task, was represented in neural activity. By combining behavioral, subjective, and neural evidence, the present study sought to clarify both the nature of confidence and its neural representation.

In summary, the present study addressed three interrelated questions. First, how can we validate a measure of confidence? Second, is confidence better understood as a causal determinant of behavior or as a state generated by environmental contingencies? And third, is information about confidence represented in neural activities measured with EEG and fMRI? To address these questions, we conducted complementary behavioral and neuroimaging experiments based on the same decision-making task.

## Results

### Experiment 1

Across two days, sixteen participants performed a decision-making task (Fig. 1a). A pacman-like stimulus (*pacman*, hereafter) was presented with two different fruits (e.g., cherry and banana) and participants had to choose the pacman’s favorite fruit based on its appearance^26^. Three visual features constituted the pacman: color (blue/yellow), line orientation (horizontal/vertical), and mouth direction (left/right). Two of the three features determined the fruit preference, the combination of which is hereafter referred to as a *feature rule*, with the third one being irrelevant. Hence, there were three possible feature rules: color-line, color-mouth, and line-mouth. Participants had to guess which rule was in effect based on the feedback for fruit choices (point gains/losses), like the Wisconsin Card Sorting Test ^27^.

**Fig. 1:**
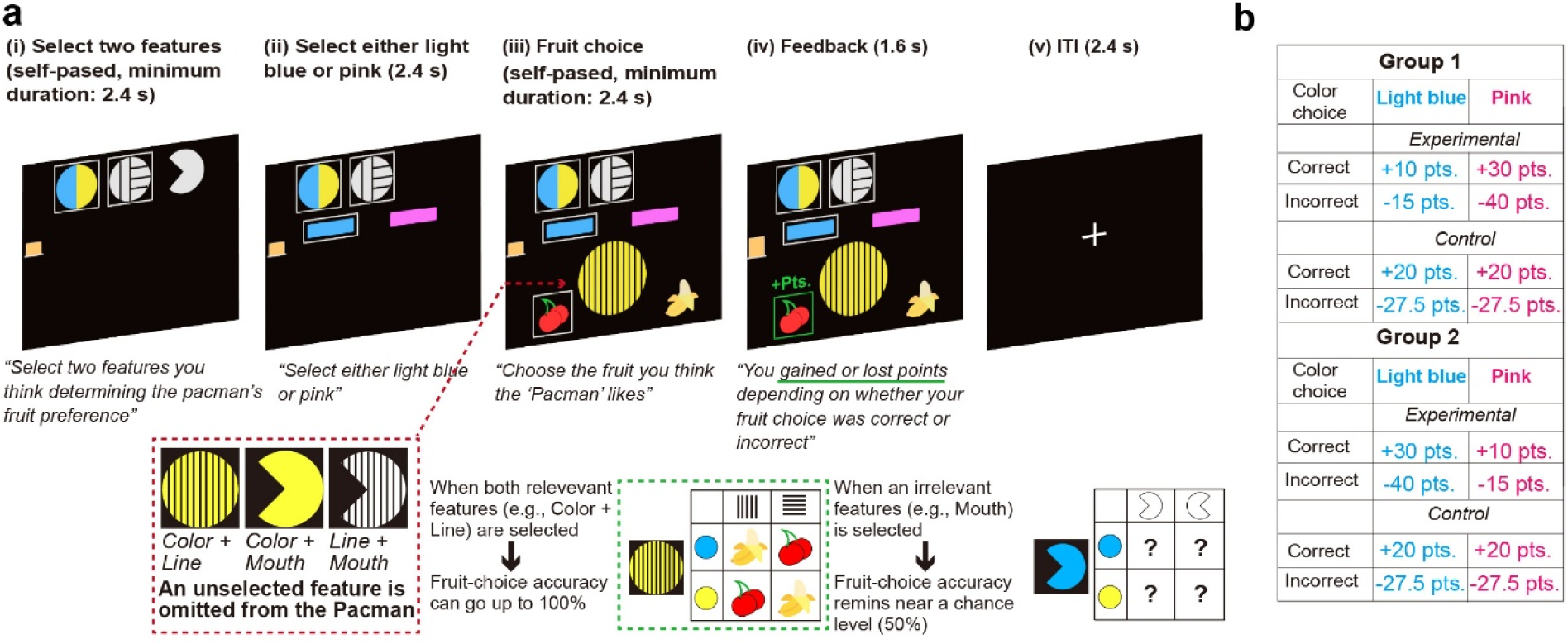
An overview of the decision-making task in Experiment 1. **a,** Task design. Participants inferred the pacman’s favorite fruit from its visual features presented on a computer monitor. The fruit preference was determined by a feature rule defined by a combination of two of the three features: color (blue/yellow), line orientation (vertical/horizontal), and mouth direction (left/right). In each trial, participants (i) selected two features, (ii) selected either the light-blue or pink option, (iii) viewed a pacman consisting of the two selected features, missing the third unselected feature, and selected a fruit, and (iv) received feedback indicating whether the fruit choice was correct (point gain) or incorrect (point loss), after which the next trial began. Participants were never verbally instructed about what the light-blue and pink options were for but could figure it out from the feedback for correct and incorrect fruit choices. For example, if color and line orientation defined the feature rule, a yellow pacman with vertical and horizontal lines might prefer cherry and banana, respectively, whereas a blue pacman with these respective lines might prefer banana and cherry. The feature rule and the fruit mapping shifted after 29.71 and 11.56 trials on average, respectively. **b,** Payoff matrix. In the experimental condition, correct and incorrect fruit choices yielded +30 and –40 pts., respectively, after selecting the pink option, and these respective fruit choices yielded +10 and –15 pts. after selecting the light-blue option for half of the participants (Group 1); functions of these colors were reversed for the other half (Group 2). This payoff matrix was to encourage selection of the high-risk options after participants made correct fruit choices across consecutive trials while encouraging the low-risk choice otherwise. In the control condition, correct and incorrect fruit choices yielded +20 and –27.5 pts., respectively, regardless of the risk choice.

Each trial consisted of the following components: i) selection of two features, ii) a choice between two colors (light-blue/pink), iii) pacman presentation and fruit choice, iv) fruit-choice feedback, and v) intertrial interval (ITI). The feature selection affected how the pacman would look: The pacman included only the two selected features (e.g., color-line), missing the third unselected feature (e.g., mouth). Importantly, participants could gain points as long as their fruit choices were correct despite selecting an irrelevant feature (e.g., mouth). In that case, however, it was difficult to keep making correct fruit choices because one of the relevant features (e.g., line) was missing from the pacman. Also important was that the choice between light-blue and pink adjusted point gain/loss amounts for fruit choices according to a payoff matrix (Fig. 1b). The payoff matrix differed between experimental and control conditions. In the experimental condition, correct and incorrect fruit choices after selecting a color (e.g., light-blue) led to +30 and –40 points, respectively, whereas those respective choices after selecting the other color (e.g., pink) led to +10 and –15 points. In this payoff matrix, the former color served as a high-risk/high-return choice (*high-risk choice*, hereafter) and the latter as a low-risk/low-return (*low-risk choice*, hereafter) choice. In the control condition, these risk choices led to identical outcomes/returns: Correct and incorrect fruit choices resulted in +20 and – 27.5 points, respectively. The feature rule and the mapping for fruit preference (*fruit mapping*, hereafter; e.g., a yellow pacman with vertical lines preferred cherry) occasionally shifted throughout the task. Each condition had six consecutive blocks of 54 trials (324 trials total).

For the analysis, we first examined whether participants performed the decision-making task as we intended. Fig. 2a shows time courses of feature-choice accuracy, fruit-choice accuracy, and high-risk choice for a representative participant, with each measure being the moving average of five trials (see Fig. S1 for an entire set of participants included in the analysis). For this and subsequent analyses, correct feature choice, correct fruit choice, and high-risk choice were coded as 1 and as 0 otherwise. General trends of these three measures were similar to each other in the experimental condition. In contrast, their trends considerably differed, especially between fruit-choice accuracy and high-risk choice in the control condition. The latter result was expected given no functional difference between high- and low-risk choices in that condition. To further examine this, Fig. 2b shows coefficients of cross-correlations between fruit-choice accuracy and high-risk choice, averaged across 16 evaluable participants (see Method for exclusion criteria). The coefficient was highest at lags of +1 and +2 trials in the experimental condition, indicating fruit-choice accuracy changed first, followed by changes in risk choices. In contrast, the cross-correlation coefficients remained near zero in the control condition. Fig. 2c shows high-risk choice, fruit-choice accuracy, and feature-choice accuracy before and after shifts in feature rule and fruit mapping, averaged across the 16 participants. The high-risk choice (left panel) in the experimental condition was relatively high before those shifts and then decreased to low levels afterwards, whereas this measure changed little in the control condition. In contrast, fruit-choice and feature-choice accuracies (middle and right panels) differed little between experimental and control conditions, indicating task performance in terms of accuracy was little disrupted by making the outcomes of high- and low-risk choices identical. Overall, these behavioral measures indicate that these participants performed the task as we intended (see Fig. S2 for other participants whose task performance met our exclusion criteria).

**Fig. 2:**
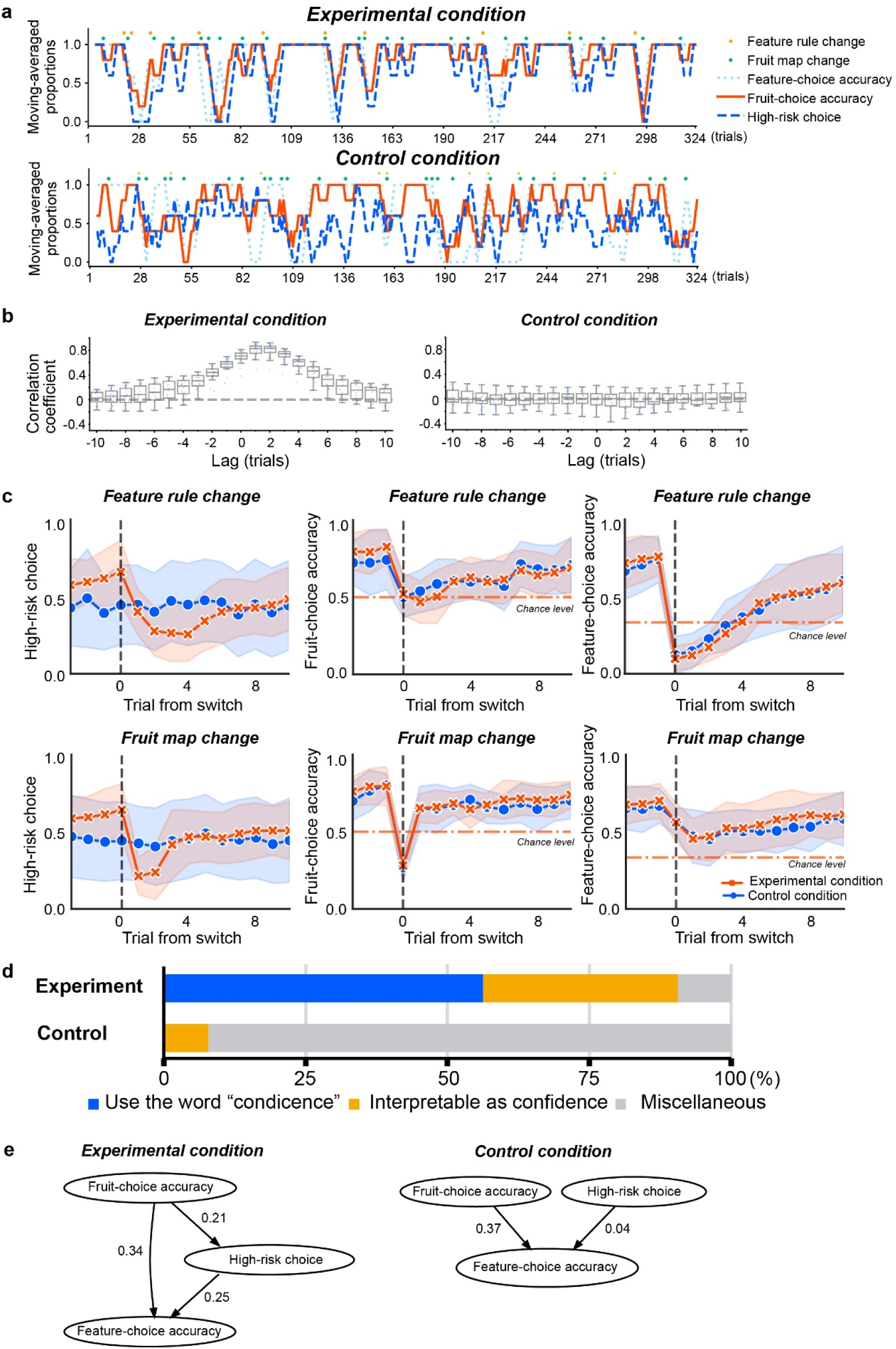
Results in Experiment 1. **a,** Time courses of feature-choice accuracy, fruit-choice accuracy, and high-risk choice, smoothed using a 5-trial moving average, for a representative participant. Correct feature choice, correct fruit choice, and high-risk choice were coded as 1 and as 0 otherwise. The upper and lower panels are for the experimental and control conditions, respectively. During the task, feature rules and fruit mappings occasionally shifted, independently of each other. **b,** Coefficients of cross-correlations between high-risk choice and fruit-choice accuracy in the experimental (left panel) and control (right panel) conditions. The box plot at each trial lag represents a summary for 16 evaluable participants. The coefficients were computed for lags ranging from −10 to +10 trials, using the moving average of each measure with a 5-trial window. **c,** High-risk choice, fruit-choice accuracy, and feature-choice accuracy around switches (shifts) in feature rule and fruit mapping. Trial 0 on the *x*-axis corresponds to the trial in which the feature rule or the fruit mapping shifted. The plot shows averages across 16 evaluable participants in a range from −3 to +10 trials relative to Trial 0. **d,** A summary of the classification of participants’ questionnaire answers in the experimental (top panel) and control (bottom panel) conditions. The blue bar represents those who directly used words like confidence and certainty/uncertainty, the orange bar for those whose descriptions of their feelings were interpretable as expressions of confidence, and the gray bar for miscellaneous answers. **e,** Direct acyclic graphs showing the inferred causal structure among feature-choice accuracy, fruit-choice accuracy, and high-risk choice in the experimental (left panel) and control (right panel) conditions, estimated using DirectLiNGAM.

After each condition, participants completed a questionnaire, which had the following open-ended questions:

Tell us about situations in which you chose the light blue: a) During the task, what feeling did you have when selecting this color?; and b) What was the situation like when you selected it?

The questionnaire had the same set of questions for the pink option as well. These questions were to examine a participant’s subjective experience and situations associated with the subjective experience. We were particularly interested in whether they would describe their feelings in a way interpretable as certainty/uncertainty in the experimental condition, but not in the control condition. Importantly, in both conditions, we neither used the words “confidence”, “certainty”, “uncertainty”, and the like nor provided verbal instructions about values in payoff matrices throughout the experiment. Four raters (two co-authors and two graduate students who were not informed about the present experiment, all native Japanese speakers) independently classified participants’ questionnaire answers (see Table 1 for samples) into the following categories: A) Participants directly used words like “confidence” and “certainty”, or antonyms of these words; B) they did not use those words but descriptions of their feelings were interpretable as such; and C) neither/miscellaneous.

**Table 1.** Examples of participants’ answers to the post-task questionnaire. Words like *confidence* are highlighted in red. The questionnaire asked participants to describe their subjective experience and situations associated with the subjective experience in an open-ended manner. All written responses were in Japanese and were then translated into English, as shown here, using OpenAI’s ChatGPT with the prompt “Translate this text into English” to minimize a human translator’s bias.

|  |  | Control |  | Experimental |  |
|---|---|---|---|---|---|
| ID |  | “Low risk” | “High risk” | “Low risk” | “High risk” |
| 2 | Subjective experience | A) Mainly during the first experiment and throughout the sessions. | A) Throughout the experiment | I chose it when I wasn’t **confident** about the rule. | I made my choice when I was **confident** about the rule. |
|  | situation | B) Because it’s my favorite color. | B) No particular awareness. | At the beginning of the questions, I felt that I couldn’t figure out the rule. | I felt I understood the rule’s logic. |
| 9 | Subjective experience | I didn’t really care much about it. | I didn’t really care much about it. | When I didn’t know the correct answer. | When I was **confident** in the correct answer. |
|  | situation | I didn’t really care much about it. | I didn’t pay particular attention. | When the features had changed | When I was in a situation of giving consecutive correct answers |

Fig. 2d shows the mean percentage of the participants classified into each category, averaged across the four raters (Inter-rater agreement was high with Fleiss’s κ=.82; see Table S1 for an entire set of questionnaire answers). Notably, the percentages were 56.25, 34.38, and 9.37% for Categories A, B, and C, respectively, in the experimental condition. In contrast, the respective percentages were 0.00, 7.81, and 92.19% in the control condition. Thus, participants were much more likely to report the feeling of confidence and the like when high- and low-risk choices led to differential outcomes than when those led to identical outcomes. To examine this statistically, we conducted a generalized linear mixed model (GLMM) with a binomial error distribution and a logit link function, using the *fitglme* function in MATLAB with the following arguments: data, ‘y ∼ 1 + Condition + (1|Subject) + (1|Rater)’, ‘Distribution’, ‘Binomial’, ‘Link’, ‘logit’. Specifically, Categories A and B were grouped together as indicating the presence of confidence, whereas Category C was considered to indicate the absence of such a feeling. This binary variable served as the dependent variable in the GLMM. The experimental condition was associated with a higher probability of feeling confidence than the control condition, *β* = 5.97, *SE* = 0.85, *t* = 7.05, *p* = 1.07e^-10^.

To further analyze the relation between high-risk choice and other task-performance variables, we conducted generalized estimating equation (GEE) analyses using feature-choice accuracy and fruit-choice accuracy as predictors of high-risk choice (see Method). Both measures significantly predicted high-risk choice in the experimental condition, but fruit-choice accuracy showed the strongest effect when entered simultaneously with feature-choice accuracy (Table 2). This suggests that high-risk choice was most strongly associated with the accumulated history of making correct fruit choices. To visually characterize the structure of these relations, directed acyclic graphs (DAGs) were estimated separately for the experimental and control conditions using DirectLiNGAM, a causal discovery algorithm (see Method). In the experimental condition, fruit-choice accuracy influenced *both* high-risk choice and feature-choice accuracy (Fig. 2e). In addition, high-risk choice mediated the link between fruit-choice accuracy and feature-choice accuracy. The latter result suggests that making correct fruit choices led to a higher probability of selecting a high-risk choice, which was more likely to be followed by the selection of correct feature rules. In contrast, the mediation by high-risk choice disappeared in the control condition: Fruit-choice accuracy only influenced feature-choice accuracy but not high-risk choice whereas high-risk choice little influenced feature-choice accuracy. Thus, just by looking at the result in the experimental condition, confidence (indicated by high-risk choice) seemingly influenced behavior as a mediator when risk choices led to differential outcomes. Such mediation might promote the view that confidence causally affects behavior. Nonetheless, the mediation disappeared when the risk choices led to identical outcomes in the control condition, suggesting that the mediation itself is a product of the environmental contingency.

**Table 2.**
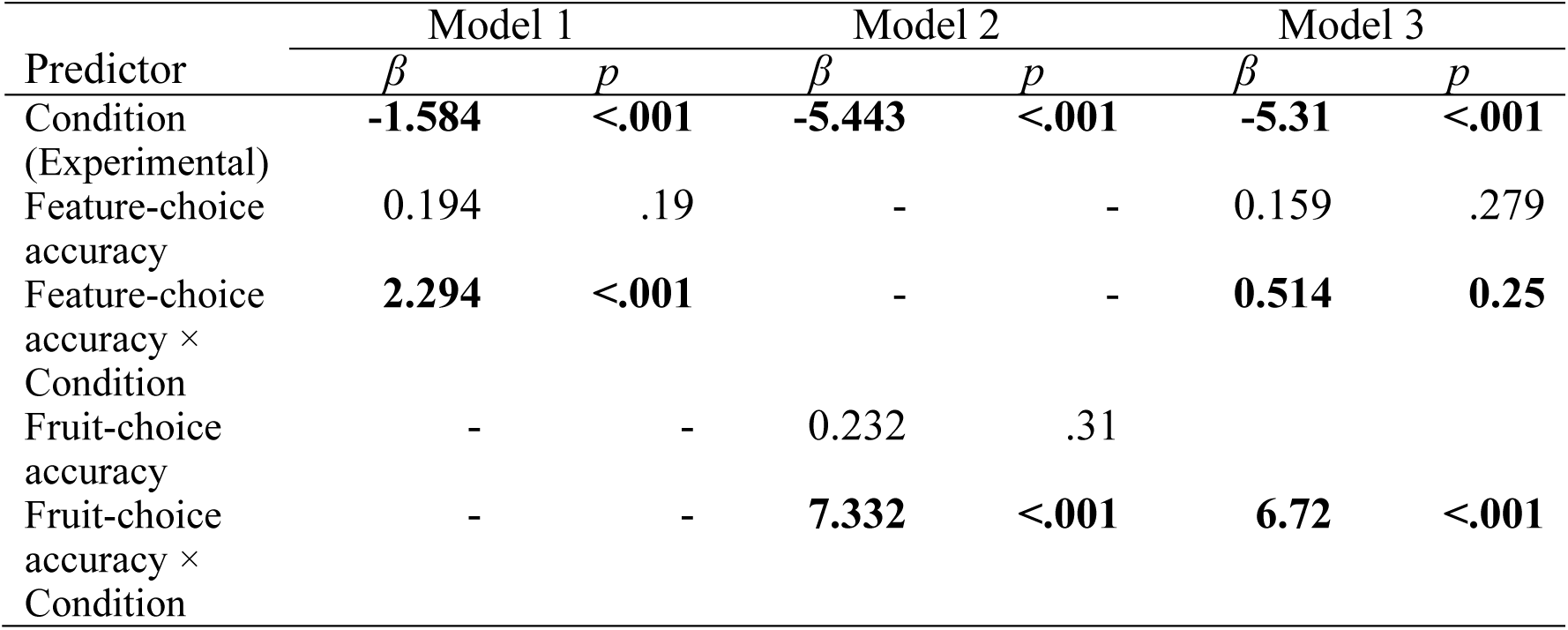
A summary of generalized estimating equation (GEE) analyses examining whether task-performance variables predict high-risk choice. Model 1 included feature-choice accuracy, Model 2 included fruit-choice accuracy, and Model 3 included both of the predictors. Regression coefficients (*β*) and *p*-values are reported. Significant interactions with condition indicate that associations between task-performance variables and high-risk choice were selectively enhanced when high-and low-risk choices led to differential outcomes.

|  | Model 1 |  | Model 2 |  | Model 3 |  |
|---|---|---|---|---|---|---|
| Predictor | *β* | *p* | *β* | *p* | *β* | *p* |
| Condition (Experimental) | **-1.584** | **<.001** | **-5.443** | **<.001** | **-5.31** | **<.001** |
| Feature-choice accuracy | 0.194 | .19 | - | - | 0.159 | .279 |
| Feature-choice accuracy × Condition | **2.294** | **<.001** | - | - | **0.514** | **0.25** |
| Fruit-choice accuracy | - | - | 0.232 | .31 |  |  |
| Fruit-choice accuracy × Condition | - | - | **7.332** | **<.001** | **6.72** | **<.001** |

Overall, the participants felt certain and uncertain when high-and low-risk choices led to differential outcomes and such feelings disappeared when these risk choices led to identical outcomes. This indicates that confidence is a *product* of contingencies between choices and their differential outcomes, supporting the notion that confidence is an organism’s state controlled by environmental contingencies (an intervening variable) rather than a cause of behavior (a hypothetical construct). Otherwise, the cause should not disappear with environmental manipulations if confidence affects behavior independently from those manipulations. This was not apparent in previous studies on confidence presumably because a contingency typically remains present throughout an experiment^4^; thus, feelings of certainty and uncertainty also remain, which may bolster the notion that those feelings causally affect behavior. When the contingency is removed as done in the present experiment, however, those feelings likely disappear eventually.

The present results also indicate that a participant’s choice between high- and low-risk options in the present decision-making task can safely be regarded as a behavioral measure of confidence when those choices lead to differential outcomes (assuming that task performance does not meet our exclusion criteria; see Method). We took advantage of this in the search for neural correlates of confidence in the next experiment.

### Experiment 2

We examined whether we could identify neural correlates of confidence and decode it from simultaneous fMRI-EEG data acquired during a decision-making task. The task was similar to the experimental condition in Experiment 1 except that we explicitly added the labels “High” and “Low” to high- and low-risk options, respectively, and there was no post-task questionnaire. These procedural modifications were based on the consideration that simultaneous recording of fMRI and EEG was so costly that having a control condition was not a practical option. Moreover, the contingency between risk choices and their differential outcomes was responsible for the feelings of certainty/uncertainty in Experiment 1, suggesting that providing the explicit labels would not hurt the interpretation of data. Data from 41 evaluable participants, different from those who served in Experiment 1 (except for 7 participants; see Method), were analyzed. Noise in EEG arising from fMRI recording was removed (Fig. S4) and the task performance of the 41 participants was similar to that of participants in Experiment 1 (Fig. S5).

First, we investigated neural correlates of confidence using a conventional whole-brain general linear model (GLM) analysis (Fig. 3a, b; Table S2). Contrasting high-and low-confidence trials revealed significant differences in the activity of several cortical and subcortical regions. Specifically, high-confidence trials were associated with stronger activity in the supplementary motor area (SMA), middle cingulate cortex (MCC), superior frontal gyrus, bilateral insula, Rolandic operculum, superior temporal gyrus, and putamen. In contrast, low-confidence trials were associated with stronger activity in widespread frontoparietal regions, including the inferior parietal lobule (IPL), angular gyrus, superior parietal lobule (SPL), middle frontal gyrus (MFG), superior frontal gyrus (SFG), precuneus, and caudate nucleus. These suggest that confidence is associated not with a specific area of the brain but rather with distributed areas across frontal, parietal, insular, and cingulate networks. This finding may not be surprising given confidence is part of metacognition, involving other cognitive functions.

**Fig. 3:**
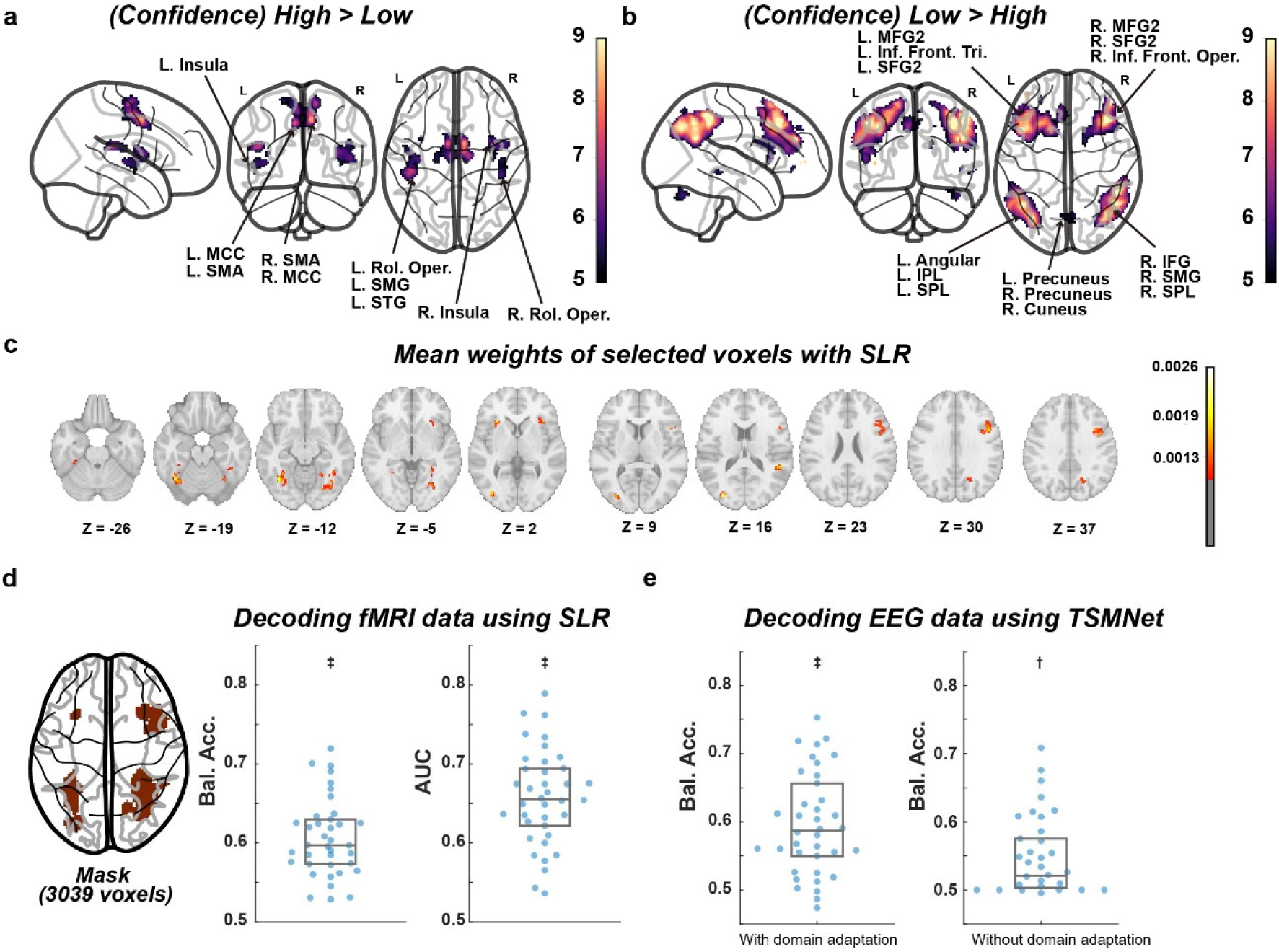
Neural correlates and decoding of confidence in Experiment 2. **a,** Whole-brain GLM contrast of high-confidence > low confidence. **b,** Whole-brain GLM contrast of low confidence > high confidence. **c,** Mean absolute weights of voxels selected by sparse logistic regression (SLR). **d,** Decoding accuracy of confidence from fMRI activity using the SLR. The left panel shows a spatial mask (3039 voxels) derived from a previous study (Cortese et al., 2021). The middle panel shows balanced accuracies obtained with leave-one-subject-out (LOSO) cross-validation, and the right panel shows the corresponding area under the receiver operating characteristic curve (AUC, ‡p < 1e-5). **e,** Decoding of confidence from EEG signals using a linear classifier with the softmax function with LOSO cross-validation after applying Tangent Space Mapping Network (TSMNet; Kobler et al., 2022) to preprocessed EEG. The left and right panels show balanced accuracy obtained with and without domain adaptation (i.e., geometrical alignment for inter-subject differences), respectively, within the TSMNet framework (†p < 1e-4, ‡p < 1e-5).

We next tested whether trial-by-trial confidence levels could be decoded from fMRI activity patterns (see the “Decoding high/low confidence from fMRI data” subsection in Method). This analysis was to determine whether confidence-related information was represented in fMRI activity patterns at the single-trial level. Using a spatial mask relevant to confidence derived from a previous study using a similar decision-making task^26^ (the left panel of Fig. 3d) and sparse logistic regression (SLR)^28^, we trained a decoder with leave-one-subject-out (LOSO) cross-validation. The decoder classified high- and low-confidence trials above chance level, achieving a balanced accuracy of 0.61 ± 0.04 (*p* = 7.6e-16, *t* = 13.47) and an area under the curve (AUC) of 0.66 ± 0.06 (*p* = 2.5e-18, *t* = 16.12) using *t*-tests (the middle panel of Fig. 3d). To identify regions contributing substantially to decoding, we examined the voxels selected by SLR. The highest contributions were observed at the inferior frontal gyrus, insula, fusiform gyrus, superior temporal gyrus, lingual gyrus, and occipital cortices (Fig. 3c; Table S3). These findings suggest that confidence-related information was broadly distributed across frontal, insular, and occipitotemporal regions, corroborating the above GLM results.

Finally, we examined whether trial-by-trial confidence levels could also be decoded from EEG signals. We trained a linear classifier with the softmax function with LOSO cross-validation after applying Tangent Space Mapping Network (TSMNet)^29^ to preprocessed EEG (see the “Decoding confidence from EEG” subsection in Method). The classifier successfully discriminated high- and low-confidence trials with a balanced accuracy of 0.55 ± 0.05 (*p* = 1.2e-5, *t* = 5.04), significantly above chance level (the right panel of Fig. 3e). Applying a domain adaptation method in order to geometrically align inter-subject differences in the TSMNet framework further improved decoding performance to a balanced accuracy of 0.60 ± 0.07 (*p* = 1.1e-9, *t* = 8.04; the left panel of Fig. 3e). These results indicate that confidence-related information also was represented in EEG signals.

Taken together, these results demonstrate that confidence can be reliably decoded across individuals from both fMRI activity patterns and EEG signals. The findings indicate confidence-related information is represented in neural activity and can be detected across different neuroimaging modalities.

## Discussion

Confidence is a widely used concept, yet its relation to behavior remains unclear, particularly regarding whether it acts as a causal determinant of behavior or merely reflects an organism’s state controlled by environmental variables. To study confidence, researchers have typically relied on self-ratings by participants or behavioral measures operationally defining confidence. However, these approaches face a common limitation: Confidence is not directly observable by a third party, making measurement validation and analyses difficult. Moreover, some researchers claim that they identified neural correlates of confidence, but it is a logical fallacy to treat these correlates as evidence for confidence because that itself is operationally defined. To tackle these issues, we developed a procedure combining a decision-making task, risk choices, and open-ended questions in a post-task questionnaire. In Experiment 1, many participants felt certain and uncertain when making high- and low-risk choices, respectively, during the task as evident in their questionnaire answers. These feelings almost disappeared when the risk choices led to identical outcomes, suggesting confidence is a state controlled by environmental contingencies rather than a cause of behavior. In Experiment 2, we demonstrated that confidence measured with the validated method was represented in neural activities (fMRI and EEG) without falling into the above logical fallacy. In what follows, we discuss each of these points in more detail.

The present study provides one of the clearest demonstrations of the validity of a confidence measure thus far. Self-ratings are frequently used to measure confidence, often through explicit instructions asking participants to rate the level of confidence about their decision ^30^, but there is no direct way to verify whether these ratings accurately reflect what they actually feel. In other studies, particularly those involving nonhuman animals, researchers have operationally defined confidence using behavioral measures such as reaction time, wagering, or choice^16^. The operational definition alone does not provide evidence that those behavioral measures indeed reflect confidence (tautological, otherwise). Neural correlates provide another avenue for examining confidence. Yet their validity ultimately depends on the validity of the underlying behavioral measures^24, 31^. One important precedent was the work of Smith et al. (1995)^32^, who asked participants during post-experimental interviews to describe their feelings when selecting an uncertain option. However, details of the interview were not provided.

The present procedure extends previous wagering-type approaches to confidence measurement^10^. First, we introduced a payoff matrix designed to minimize exclusive choices for a particular risk option, an issue previously discussed by Clifford et al. (2008)^33^. The present payoff matrix promoted high- and low-risk choices when decision accuracy was high and low, respectively. Second, we combined the risk choices with an open-ended post-task questionnaire, allowing us to directly evaluate whether participants experienced feelings of certainty/uncertainty. Importantly, in Experiment 1, we gave participants no instruction about confidence (and no instruction about how many points they would gain/lose) throughout the experiment. Nevertheless, approximately 90% of evaluable participants reported feelings interpretable as certainty/uncertainty. These reports therefore provide strong support for the present procedure capturing the subjective experience of confidence.

The central question in the present study was whether confidence should be regarded as a cause of behavior (i.e., a hypothetical construct that can causally affect behavior beyond the effects of observable events) or as an organism’s state controlled by environmental variables (i.e., an intervening variable which summarizes the relation among observable events without independent causal status). Our results support the latter interpretation. Confidence reports were strongly associated with the contingency between choices and their differential outcomes. When this contingency was present, 90% of participants reported feelings consistent with certainty/uncertainty; when it was removed, such reports largely disappeared in the control condition (only 8% of participants reported such feelings). Because confidence itself almost disappeared in the absence of the contingency with all other aspects of the experimental procedure held constant, it becomes difficult to maintain the common assumption that confidence causally affects behavior^6,7,8,9^. Instead, our findings suggest that confidence is better understood as a *product* of environmental contingencies rather than a causal agent driving behavior. Although the findings do not rule out such a causal influence under other conditions (it is difficult to prove a negative), they demonstrate that confidence and behavior can systematically covary in response to experimentally manipulated environmental contingencies. In contrast, we are not aware of any direct evidence supporting the claim that confidence causally affects behavior beyond the effects of observable events.

The present findings also raise the question of why confidence is often interpreted as a causal determinant of behavior. One reason may be that confidence and choice typically occur together^16^. High confidence often accompanies successful choices and low confidence accompanies a mixture of successful and unsuccessful choices, making it intuitive to attribute behavioral changes to confidence rather than to the environmental contingency that simultaneously affects both confidence and choice (i.e., fundamental attribution error) ^34^. Furthermore, experiences accumulated over a lifetime may contribute to feelings of certainty/uncertainty even when explicit contingencies are absent in a particular situation. Such a history effect may persist for some time (which may explain why 8% of our participants reported feelings interpretable as certainty/uncertainty in the control condition) but may eventually disappear if the contingency remains absent for a sufficiently long time. Together, these factors may foster the impression that confidence independently causes behavior, whereas the present results suggest that confidence is more appropriately viewed as a state arising from interactions between an individual’s choice and differential outcomes from the environment. The mechanism underlying the latter interpretation (i.e., selection by consequences) is conceptually similar to Darwin’s theory of evolution by natural selection^35^.

As for neural correlates of confidence, we decoded confidence-related information from both EEG and fMRI signals in Experiment 2. Trial-by-trial confidence levels were successfully classified from distributed fMRI activity patterns across participants, indicating that confidence-related information was represented in neural activities at the single-trial level (see also Fig. S6 for the result of an additional fMRI analysis with a hidden Markov model, identifying several brain states associated with high and low confidence in a data-driven manner). We also decoded confidence from simultaneously recorded EEG signals, demonstrating that confidence-related information was also accessible from scalp-recorded EEG activities. Notably, EEG decoding performance further improved when a domain-adaptation procedure (i.e., a geometric transformation for reducing inter-subject variability in data) was applied. Together with the decoding results, these findings suggest that confidence-related information is reflected not only in static neural representations but also in dynamic fluctuations of brain states. Importantly, our neuroimaging analyses were not intended to identify neural correlates of confidence operationally defined using behavioral indices, as was done in previous studies^17,21^. Rather, we aimed to determine whether confidence measured using a validated method was reflected in neural activities. This represents an advancement toward better understanding of the relation among behavior, subjective experience, and the brain.

Several limitations should be addressed. First, a substantial proportion of recruited participants were excluded because they failed to achieve sufficient task performance or showed little sensitivity to reward/loss contingencies (see Methods for details^36,37,38^). This suggests that the current task may have been overly difficult for some individuals. Future studies may improve participant retention by reducing task complexity, reducing the probability of feature-rule/mapping shifts, or adjusting the payoff matrix. Second, reporting confidence-related feelings in the post-task questionnaire did not necessarily mean that participants felt certain and uncertain every time they selected high- and low-risk choices, respectively. Participants likely needed some experience with the payoff matrix before those feelings emerged. Therefore, the questionnaire answers should be treated as evidence supporting the validity of the present task for measuring confidence rather than the ground truth of certainty/uncertainty in every single trial. Third, unlike in Experiment 1, we explicitly instructed participants to report whether they were certain or uncertain when making a choice between high-and low-risk options in Experiment 2. We instead assumed that the contingency between risk choices and differential outcomes would generate those feelings. Nonetheless, evidence for neural correlates of confidence would have been more convincing if the explicit instruction had been absent.

In conclusion, we developed a novel procedure for measuring confidence that incorporated a step for measurement validation. In this procedure, participants reported feelings of certainty/uncertainty when risk choices led to differential outcomes, but those feelings almost disappeared when the contingency was absent. These results support the interpretation of confidence as an organism’s state (an intervening variable) rather than a cause of behavior (a hypothetical construct). We further demonstrated that both EEG and fMRI signals contained information sufficient to discriminate between high and low confidence. Together, these findings challenge conventional interpretations of confidence as a causal determinant of behavior and establish an experimentally grounded framework for distinguishing the environmental determinants, behavioral correlates, and neural representations of subjective confidence. More broadly, the present study addresses a fundamental challenge in psychology and neuroscience: how to investigate the nature and neural basis of subjective experiences without conflating correlation with causation.

## Methods

The protocol of this study was approved by the ATR Review Board Ethics Committee (IRB committee number:18000062; application number: 167) and followed the Declaration of Helsinki. All participants provided written informed consent prior to their participation with all instructions provided in Japanese. As part of the consent process, participants explicitly agreed to the collection of demographic information (age and sex), behavioral task data, and neural activity data. All data were anonymized to protect participants’ privacy.

### Experiment1

#### Participants

Initially, 29 participants (19 men, 10 women) were recruited through public advertisements in the local community with the following criteria: 1) ages ranging from 20 to 49; and 2) no color blindness. These participants were quasi-randomly assigned to one of four sub-groups (for counterbalancing the order of experimental and control conditions and the risk-choice colors) with the restriction that the sub-groups would have equal numbers of participants. To be included in the final analysis, however, a participant’s task performance had to meet criteria described below. Thirteen did not meet the criteria, resulting in a final set of 16 participants (9 males, 7 females; mean age = 23.4 years, SD = 1.93) with each sub-group having 4 participants. Mean values shown in the main results were computed across these 16 participants while Figs. S2 and S3 show data for the excluded participants.

#### Apparatus

The experiment was conducted in a small room which had a table, a chair, ceiling lights, and an air conditioner. On the table were a conventional numeric keypad (REALFORCE) and a computer monitor (BenQ®; 60 cm by 34 cm; a 1920x1080 resolution) with its center approximately at eye level when participants sat on the chair. We used only three keys on the keypad: “4”, “8”, and “6” keys which served as “left”, “center”, and “right” keys, respectively. Presses on other keys had no programmed consequence and were not recorded. The monitor was connected to a Windows 10 computer with cables located in an adjacent room. A custom program written in Python (ver. 3.8.18) running on the computer controlled and recorded experimental events.

#### Procedure

Each participant had two 3-hr laboratory visits, separated by a range of 1 to 41 days (mean=11.5 days). One visit was for an experimental condition and the other for a control condition, with the condition order counterbalanced across participants. Each laboratory visit consisted of a decision-making task followed by a post-task questionnaire. In that task, 1 point was worth 1 JPY and a payment amount was computed based on task performance in each block of 54 trials, rounded to the nearest multiple of 50 JPY (e.g., points in a range from 25 to 74 were rounded to 50 JPY). Negative points in a block were treated as 0 JPY. Payment was made at the end of each laboratory visit, and the payment included the total payment across blocks (with a maximum of 5,000 JPY) and base pay for participation (6,000 JPY). Answers on the questionnaire did not influence the payment amount. The mean task-performance payment for the two laboratory visits was 2776.56 JPY for the 16 evaluable participants.

Each condition had six main blocks of 54 trials, totaling 324 trials. Each trial consisted of four components in the following order: i) selection of two features, ii) a choice between two colors (light-blue/pink), iii) pacman presentation and fruit choice, and iv) fruit-choice feedback (see Fig. 1a). The feature-choice and fruit-choice components lasted for at least 2.4 s and continued until participants made a choice. In the latter case, the Python program checked whether a choice was made at 0.8-s intervals and proceeded to the next component upon the detection of a choice (The 0.8-s interval in this and all other components of a trial was to be consistent with how experimental events occurred during an fMRI recording in Experiment 2). The risk-choice component ended as soon as 2.4 s elapsed since its onset. Likewise, the fruit-choice feedback component ended in 1.6 s, followed by an ITI of 2.4 s.

The participant’s task was to infer the fruit preference (e.g., cherry v. banana) of a pacman-like stimulus (*pacman*, hereafter) presented on the monitor based on its appearance ^26^. Three visual features, with each having binary parameters, constituted the pacman: color (blue/yellow), line orientation (horizontal/vertical), and mouth direction (left/right). In the feature-choice component, three icons representing these features were presented on the top side of the monitor. (The order of the icons remained the same throughout a block of 54 trials but was shuffled at a block transition.) Using the left, center, and right keys, participants selected two of the three features that might constitute a rule governing the pacman’s fruit preference. A surrounding white box appeared when a feature icon was selected and disappeared when selected again. Suppose a combination of color and line orientation constituted the rule. In this case, a yellow pacman with vertical and horizontal lines may prefer cherry and banana, respectively; a blue pacman with those respective lines may prefer banana and cherry; and mouth directions of the pacman would be irrelevant to the fruit preference (see Fig. 1a). A combination of two features governing the pacman’s fruit preference will hereafter be called a *feature rule*.

Next, in the risk-choice component, two buttons (light-blue and pink) appeared just below the feature icons. Their positions (left or right) were randomized across trials. At the component onset, one of the buttons was randomly selected automatically, indicated by a white box surrounding the button. Using the left and right keys, participants could switch it to the other button if they wished; otherwise, they did not have to press a key. The color having the white box at the end of the component was considered as a final choice. The choice between two colors determined the point-gain amount for correct fruit choices and the point-loss amount for incorrect fruit choices in the same trial, and this also depended on the condition (see Fig. 1b). The payoff matrix in an experimental condition was as follows: The light-blue option followed by correct and incorrect fruit choices led to +10 and −15 pts., respectively; the pink option followed by these respective fruit choices led to +30 and −40 pts. In this case, the light-blue served as a low-risk/low-return choice (*low-risk choice*, hereafter) and the pink as a high-risk/high-return choice (*high-risk choice*, hereafter). The colors for high- and low-risk choices were counterbalanced across participants (see Table S1). In contrast, the payoff matrix in a control condition was as follows: Regardless of the risk choice, correct and incorrect fruit choices led to +20 and −27.5 pts., respectively. Thus, the “high-risk” and “low-risk” choices (despite being called such for the sake of simplicity) led to identical outcomes/returns.

The payoff matrix in the experimental condition was based on the consideration that participants would likely select the low-risk option if the subsequent fruit choice had only a 50% chance of being correct for minimizing the net loss (i.e., a net loss of −5 pts. for the low-risk option as opposed to a net loss of −10 pts. for the high-risk option). Conversely, they would likely select the high-risk option if the subsequent fruit choice was likely to be correct in order to gain +30 pts. for maximizing the net gain. From a subjective point of view of participants, we expected they would select the high-risk option if they were “confident” about the subsequent fruit choice and select the low-risk option otherwise. This was just our operational definition, however, and we could not confirm their subjective experiences at this point. Instead, our main interest was whether the payoff matrix in the experimental condition would generate such a subjective experience. Accordingly, the risk-choice options did not have the labels like “high confidence” and “low confidence” and we never used the words like “confidence” and “certainty”, or antonyms of these words, throughout the experiment (see pre-experimental instructions in Appendix). The payoff matrix in the control condition was for examining whether differential outcomes of the choice between the high- and low-risk options in the experimental condition were responsible for the distribution of risk choices as well as the generation of subjective experience.

In the fruit-choice component, a pacman was presented at the center of the monitor below the risk-choice options. Two fruits (e.g., cherry and banana) also were presented simultaneously on the left and right sides of the pacman. The fruit positions were randomized across trials. Participants selected one of the fruits as the pacman’s preference using the left and right keys, and the selected fruit was surrounded by a white box. They could switch the option within 2.4 s of the component onset, and the one selected at the end of the 2.4-s period was considered as a final choice. If a choice was not made within the 2.4-s period, the first choice after the period was considered as a final choice. Two aspects of the fruit-choice component were important. First, each binary parameter for a visual feature of the pacman (e.g., blue and yellow for color) had almost equal probabilities (approximately 50%) of being part of a full pacman image but its actual image only had the two features the participant had selected in the same trial, missing the third unselected feature. If participants selected color and line orientation, for instance, the pacman would have a color and lines (e.g., blue with vertical lines) but no mouth. Second, whether participants would gain or lose points in the fruit-choice feedback component was determined *solely* by whether a fruit choice was correct or incorrect, regardless of which feature rule was in effect. Given these two aspects of the fruit-choice component, participants could gain points just by chance even after selecting a feature irrelevant to the pacman’s fruit preference but it was difficult to keep gaining points across consecutive trials. Moreover, making correct fruit choices just by chance (50%) would lead to a net loss in both experimental and control conditions (see above), thereby promoting participants to search for a correct feature rule. Participants never received feedback on whether the selected feature rule was correct or incorrect – They only received feedback on whether the fruit choice was correct or incorrect, like the Wisconsin Card Sorting Test^27^. Therefore, participants had to infer the correct feature rule through trial and error, relying solely on the feedback for fruit choices.

Finally, in the fruit-choice feedback component, the white box surrounding the selected fruit turned green if the fruit choice was correct and red if incorrect and the number of points gained or lost was displayed according to the payoff matrix in effect. In addition, an orange bar located on the left side of the monitor increased/decreased by 0.54 pixels per point to indicate the current total points (each block started with an initial value of 50 pts. corresponding to a white horizontal line on the bar). This component was followed by an ITI during which a fixation cross was displayed.

To increase the task difficulty, the correct feature rule shifted to one of the other two with a 5% chance per trial but only after a minimum of four consecutive trials under the same feature rule. Moreover, the mapping for the pacman’s fruit preference (*fruit mapping*, hereafter) occasionally reversed (e.g., a blue pacman with vertical lines, which had previously preferred cherry, now preferred banana). The shift in the fruit mapping occurred with a 10% chance per trial but only after a minimum of three consecutive trials with the same preference. The feature rule and fruit mapping shifted independently from each other and stochastically during the task although the same feature rule or the same fruit mapping was selected no more than once consecutively. Each of the three possible feature rules (i.e., color/line, color/mouth, and line/mouth) had equal prior probabilities but their observed counts may not be equal. No additional balancing was imposed so that numbers of shifts in the feature rule and the fruit mapping varied across participants. Both the feature rule and the fruit mapping in the last trial of a preceding block were carried over to the first trial of the upcoming block although shifts in these could coincide with a block transition.

Before the first main block in each laboratory visit, participants completed three structured practice blocks. The practice was to familiarize participants with the task with minimal verbal instructions, especially about high- and low-risk options. Practice 1 (“forced risk choice”) was to expose participants to the outcomes of those risk options as determined by a payoff matrix. Only one option (either light-blue or pink) was presented in the risk-choice component and participants were forced to select it. They had to be exposed to all four possible combinations (i.e., light-blue followed by correct and incorrect fruit choices and likewise for pink) at least once for each combination in 20 trials; otherwise, additional trials were conducted until they were exposed to all the combinations. In this practice block, the feature rule and the fruit mapping did not shift. All other aspects of the procedure were as described for the main blocks. Practice 2 (“free choice between high and low risks”), having 20 trials, was to help participants learn how to search for a correct feature rule and a correct fruit mapping when both high- and low-risk options were available. All other aspects were as in Practice 1 except that a new pair of fruits was used, which thereafter remained unchanged until the end of a laboratory visit (i.e., different pairs of fruits were used across two laboratory visits). Practice 3 (“shifts in feature rule and fruit mapping”), having 34 trials, was to train participants to adapt to shifts in feature rule and fruit mapping. In the first trial, the feature rule and the fruit mapping were as in Practice 2. Then each shifted once at some point in subsequent trials. After completing all three practice blocks, participants proceeded to the main blocks.

The final set of a feature rule and a fruit mapping in the last trial of Practice 3 was carried over to the first trial in the first main block.

#### Exclusion Criteria

Some participants’ task performance did not meet our expectations. This included exclusive choice in the feature-choice and risk-choice components, especially in the experimental condition in which risk choices led to differential outcomes. Accordingly, we excluded participants from the analysis if they met either one of the following criteria: (a) the rate of selecting a feature rule in the experimental condition deviated by more than 20% from the rate at which the rule was correct; or (b) coefficients of cross-correlations between high-risk choice and fruit-choice accuracy at trial lags ranging from +1 to +5 were all less than 0.3 in more than two of six main blocks. These exclusion criteria were applied only to task performance in the experimental condition but not in the control condition. Participants who completed the experimental condition in the first laboratory visit and met the exclusion criteria were not invited to the second visit. As a result, 13 of 29 participants were excluded from the analysis. Data for those excluded from the analysis are reported in Figs. S2 and S3.

#### Post-task Questionnaire

Immediately after completing a condition, we handed a questionnaire to participants and instructed them, “Please fill this out”. The questionnaire consisted of the following questions: 1) the points they saw on the monitor when they made correct and incorrect fruit choices following the selection of the light-blue option; 2) the points they saw on the monitor when they made correct and incorrect fruit choices following the selection of the pink option; 3) what they felt when selecting the light-blue option, and what the situation was like when selecting it; and 4) what they felt when selecting the pink option, and what the situation was like when selecting it. The first two questions were to check to see if their description of a payoff matrix matched the one in effect in a given condition. The last two questions were to examine a participant’s subjective experiences when making risk choices. We included questions about both their feelings and situational factors because a pilot study indicated that some participants mixed up the two. If participants asked questions about the items on the questionnaire, an experimenter answered, “Write down whatever you think is right”.

#### Behavioral Modeling Using Generalized Estimation Equation Models

To analyze the relation between task-performance variables and high-risk choice, generalized estimation equation (GEE) models with a binominal error distribution and logit link function were fitted using Python using the *statsmodel* package. Specifically, GEE models were estimated using the statsmodels.formula.api.gee() function. Feature-choice accuracy and fruit-choice accuracy were entered as predictors, together with condition and their interaction terms. Participant identify was treated as the clustering variable. GEE was selected because the primary objective was to estimate population-level relations between the task-performance variables and high-risk choice while accounting for repeated observations within participants. Three models were evaluated: The first model examined the effect of feature-choice accuracy:

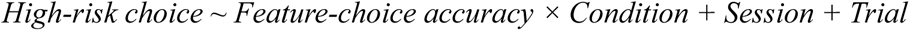

The second model examined the effect of fruit-choice accuracy:

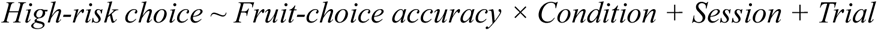

The third model simultaneously included both task-performance variables:

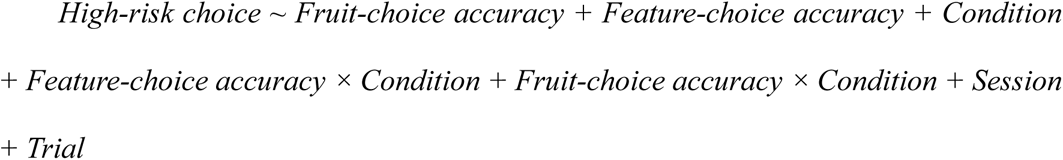

A 5-trial moving average of feature-choice accuracy and fruit-choice accuracy served as a participant’s current task-performance state. High- and low-risk choices were coded as 1 and 0, respectively.

#### Causal Structure Analysis Using DirectLiNGAM

To visually characterize the structural relations among feature-choice accuracy, fruit-choice accuracy, and high-risk choice, directed acyclic graphs (DAGs) were estimated using the DirectLiNGAM algorithm^39^ implemented in the Python package *lingam*. DirectLiNGAM is a causal discovery method for estimating directed dependencies among observed variables under the assumptions of linearity, non-Gaussian distributions, and acyclic causal structure. As in our previous application of DirectLiNGAM to fMRI data^40^, the present analysis was used to identify directed dependencies among behavioral variables and to compare network structure across conditions. Separate analyses were conducted for the experimental and control conditions. The variables entered into the analysis were feature-choice accuracy, fruit-choice accuracy, and high-risk choice (without moving average). The estimated adjacency matrices were visualized as DAGs, in which nodes represented behavioral variables and directed edges represented directional dependencies among those variables.

### Experiment 2

#### Participants

Initially, 62 participants (38 men, 24 women) were recruited through public advertisements in the local community with the following criteria: 1) ages ranging from 20 to 49; and 2) no color blindness. Participants completed a screening session before the main experiment. The screening session consisted of a decision-making task similar to the one used in the experimental condition of Experiment 1. After the screening, participants were not invited to the main experiment (1) if they met the exclusion criteria described in Experiment 1 or (2) if they took an unusually long time to complete the task. We excluded 9 participants and 1 participant for the respective criteria. In addition, 11 participants had a scheduling conflict for the main experiment; thus, they were excluded from the study. As a result, 41 participants (26 men, 15 women; mean age =23.73 years, *SD* = 3.33) proceeded to the main experiment, including 7 participants who had previously participated in Experiment 1. Unless noted otherwise, mean values shown in the main results were computed across these 41 participants.

In the main experiment, each participant had two laboratory visits (hereafter, “Day 1” and “Day 2”), separated by 1 to 51 days (mean = 8.8 days), to complete a whole set of six main blocks of 54 trials. Each laboratory visit began with EEG recording in a shield room [i.e., outside a magnetic resonance (MR) scanner]. Subsequently, simultaneous EEG-fMRI recording began within 30 min after the EEG recording. On Day 1, practice blocks described in Experiment 1 were conducted in the shielded room, and main blocks were then conducted during simultaneous EEG-fMRI. By default, participants had three main blocks on Day 1 and another three on Day 2. Due to time constraints, however, a few blocks planned for Day 1 had to be postponed to Day 2 for some participants. The payment amount for task performance in each block was computed as in Experiment 1 but with a maximum of 2,500 JPY per day. Participants received a base pay (16,000 JPY) each day in addition to the payment for task performance. The payment was made at the end of each laboratory visit. The mean of the sum of task-performance-based payments over Days 1 and 2 was 3,810 JPY.

#### Simultaneous EEG-fMRI

Details of experimental settings and data acquisitions are described elsewhere (Tsutsumi et al., 2026). Simultaneous EEG-fMRI recordings were conducted in an MR scanner. Visual stimuli were displayed on an opaque screen via a projector (Canon Power Projector, WUX6000 at a 60-Hz frame rate), being visible to participants through a small mirror attached to the head coil. A photosensor was mounted at the bottom-right corner of the screen and connected to a Stim Tracker (Cedrus® model ST-100) to record the actual time at which the stimuli were presented on the screen. Button presses were recorded using an MR-compatible response device (HHSC-1 × 4-D, Current Designs, Inc., Pennsylvania, USA).

EEG signals were recorded using MR-compatible EEG caps and amplifiers (BrainAmp MR plus, BrainAmp ExG MR, and BrainCap MR; Brain products GmbH, Germany). Each cap comprised 63 EEG channels, one electrocardiogram (ECG) channel, and four carbon-wire loop (CWL) channels. Electrode leads were connected to intermediate connecters via 30-cm cables, and the connecters were subsequently linked to the amplifiers by bundled 10-cm cables. The 63 EEG electrodes were placed on the scalp according to a modified International 10-20 system. The ground and reference electrodes were placed at AFz and FCz, respectively. The ECG electrode was positioned on the participant’s back, close to the heart. Conductive gel (Abralyt HiCl, EASYCAP, Brain Products GmbH, Germany) was used for maintaining the impedance of all electrodes below 10 kΩ throughout the experiment. Raw EEG signals were sampled at 5kHz. EEG and fMRI data were recorded simultaneously using hardware clock synchronization (SyncBox, BrainProducts, Germany). The MR scanner also sent trigger signals to mark the start time of each fMRI volume acquisition in the EEG data.

fMRI data were acquired with a multi-band sequence in a 3T MAGNETOM Prisma fit scanner, running the syngo MR VE11C/E software, with 64-channel head coils (Siemens, Erlangen, Germany). The scan protocol followed the Multimodel Harmonized MRI Imaging Protocol (HARP; see https://hbm.brainminds-beyond.jp/documents/protocol.html). All scanning parameters were as described elsewhere^41^. The brain was scanned at 0.8-s intervals (i.e., repetition time = 0.8) so that all events in the decision-making task were programmed to occur at 0.8-s intervals as well. For each of EEG and fMRI, data were preprocessed and artifacts were removed by taking steps described elsewhere^41^.

#### Decision-making task

The task was similar to the one used in the experimental condition of Experiment 1 (see Fig. 1a) with the following modifications. First, in the risk-choice component, the pink (high-risk) and the light-blue (low-risk) options had the labels “High confidence” and “Low confidence”, respectively. We included these labels to explicitly tell participants that the study was about confidence. Second, six main blocks of 54 trials were split across two days. On Day 1, participants completed practice blocks and main blocks 1–3; on Day 2, they completed main blocks 4–6. At the start of the task on Day 2, participants were instructed, “The first block today will be a continuation of the last block on Day 1.” To remind them which feature rule they selected, which fruit they chose, and whether the fruit choice was correct or incorrect in the last trial on Day 1, these images were displayed before the first trial on Day 2.

#### General linear model analysis

Statistical analyses for fMRI data were performed in a conventional way using a general linear model (GLM) implemented in SPM12 (https://www.fil.ion.ucl.ac.uk/spm/software/spm12/), following the procedures described by Ogawa et al. (2025)^42^, unless noted otherwise. Three participants were excluded from the GLM analysis (and the decoding analysis described in the next section) because at least one run contained only high-confidence or only low-confidence responses (i.e., exclusive choice), resulting in insufficient variability for reliable estimation of the corresponding GLM regressors and classifier training (see Data Sources for Fig. 3d). An event-related design was employed. Task-related regressors corresponding to trials with high and low confidence were modeled separately and convolved with the canonical hemodynamic response function (HRF). Regressors were time-locked to the onset of the risk-choice component ((ii) in Fig. S5a). To reduce potential confounds, nuisance regressors were included in the model, comprising six head motion parameters (three translations and three rotations) as well as the mean signals extracted from gray matter (GM), white matter (WM), and cerebrospinal fluid (CSF). These signals were regressed out during the model estimation. Contrast images were computed for each participant to identify brain regions associated with the trials with high and low confidence as part of the individual-level analysis (first-level analysis).

At the group level (second-level analysis), random-effects analyses were conducted to examine contrasts of “high-confidence > low-confidence” and “low-confidence > high-confidence”. Statistical analyses were restricted to a GM mask. Statistical significance was determined using a voxel-wise threshold of p < 0.001 (uncorrected), followed by cluster-wise Family-Wise Error (FWE) correction at p < 0.05. Activation maps were generated based on these thresholds.

#### Decoding high/low confidence from fMRI data

To decode confidence levels from fMRI data, we used sparse logistic regression (SLR; Yamashita et al., Neuroimage, 2008). SLR employs a hierarchical Bayesian framework with automatic relevance determination (ARD) to estimate the contribution of each voxel, enabling automatic and sparse feature selection. This approach is particularly suitable for high-dimensional fMRI data. For decoding, 3039 voxels within a predefined region-of-interest (ROI) mask^26^ were extracted. For each voxel, time series were obtained and averaged within each single trial over the time window corresponding to the risk-choice component, resulting in trial-wise activation patterns used as input features for the classifier. The classifier was trained to discriminate between high- and low-confidence trials. Model performance was evaluated using a leave-one-subject-out (LOSO; i.e., across-subject) cross-validation scheme. Because the distribution of high- and low-confidence labels varied across participants, balanced accuracy and AUC were used as performance metrics. These metrics were calculated for each cross-validation fold and subsequently assessed at the group level using one-sample t-tests to determine whether decoding performance was significantly above chance.

#### Decoding confidence from EEG

We also decoded trials with high and low confidence from EEG signals. To this end, we first examined the ratio of high- and low-confidence trials across all main blocks of trials for each participant. Of 41 participants, two had a disproportionally small number of high-confidence trials (2% and 15% respectively) compared to other participants (ranging from 37 to 76%), which would produce a decoder having a strong bias toward the high-confidence trial in the classification of EEG signals if included in decoder learning. The two participants also showed poor task performance, meeting the exclusion criteria if applied. For these reasons, we excluded these two participants from the decoding analysis. In addition, EEG preprocessing did not complete for one participant. Thus, we also excluded this participant from the analysis. Accordingly, a final dataset for EEG decoding included 38 participants (see Data Sources for Fig. 3e).

Next, we extracted 4-s epochs of EEG signals, containing the last 1.6-s segment of the feature-choice component and an entire 2.4-s period of the risk-choice component, and concatenated the epochs for each participant. A label (“high confidence” or “low confidence”) was assigned to each epoch depending on a participant’s choice in the risk-choice component. We used a conventional classifier (a linear classifier with the softmax function) for decoding after applying TSMNet^29^ to preprocessed EEG signals. Briefly, TSMNet converts a Riemannian nonlinear geometric space represented by EEG covariance matrices into a Euclidean linear space. In addition, it uses domain labels (e.g., participant) to perform domain adaptation, aligning feature distributions across domains and reducing domain-specific variability. This facilitates the learning of domain-invariant representations and improves cross-subject decoding performance. Model performance was evaluated using a LOSO cross-validation scheme.

## Acknowledgements

We thank Aurelio Cortese and Hugo Bernard Laurent Six in the Department of Decoded Neurofeedback at ATR for sharing their Python program, which served as a basis for our program for conducting the present decision-making task and also for sharing a spatial mask with us for our fMRI analysis; Miki Kitagawa for preparing for the experiment; Yoko Matsumoto for recruiting participants and scheduling the experiment. We also thank the team from the ATR Brain Activity Imaging Center (BAIC), in particular, Akikazu Nishikido, Nobuyoshi Tanki, and Akihide Yamamoto for their technical support.

## Author contributions

**Mizuki Tsutsumi**: conceptualization, methodology, writing – original draft, formal analysis

**Takeshi Ogawa**: conceptualization, methodology, writing – original draft, formal analysis

**Tomohiko Kishi**: Methodology, investigation

**Motoaki Kawanabe**: supervision, project administration, funding acquisition, writing – review & editing

**Toshikazu Kuroda**: conceptualization, methodology, software, writing – original draft

## Supplementary Materials

### Experiment 1

**Fig. S1:**
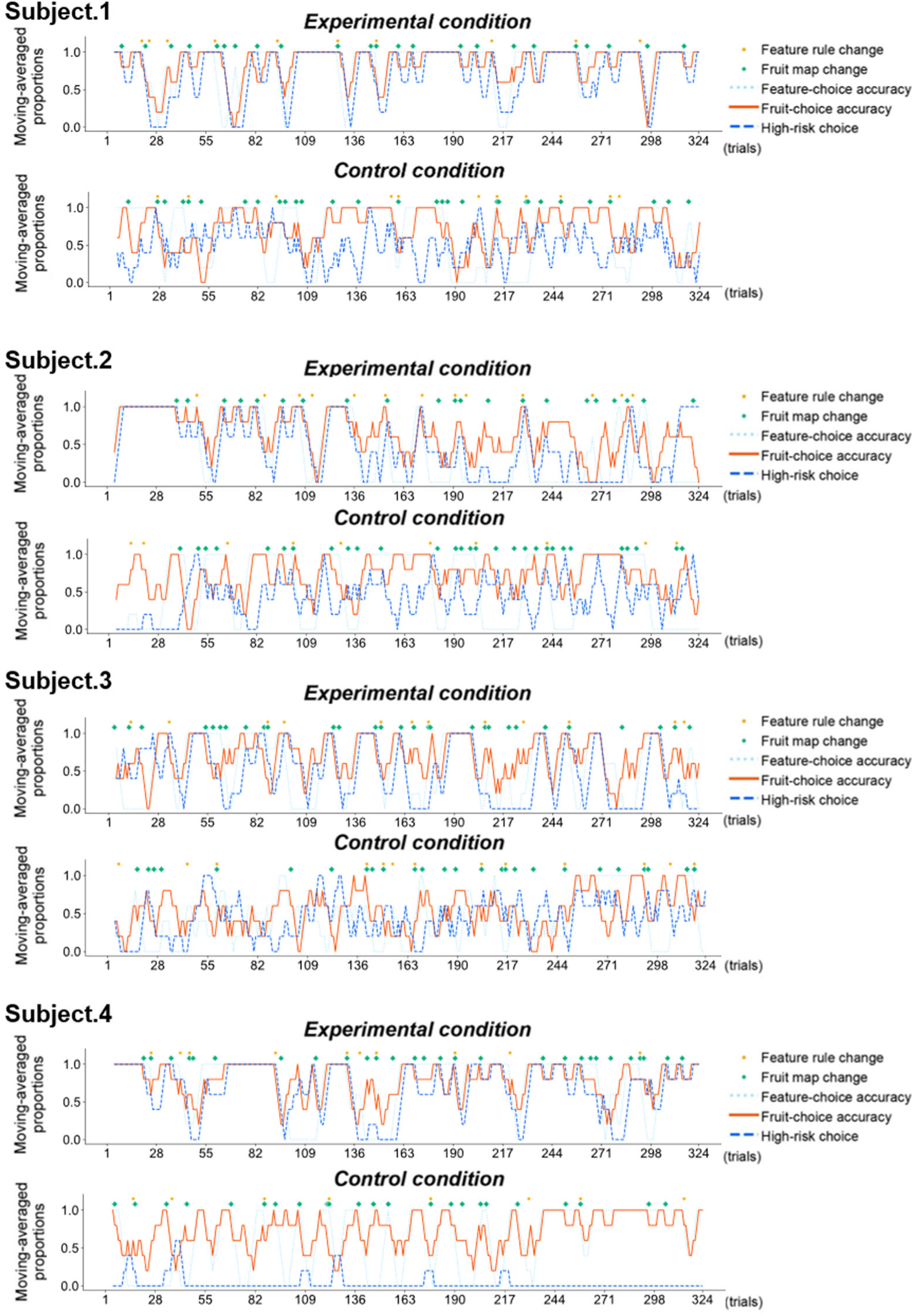
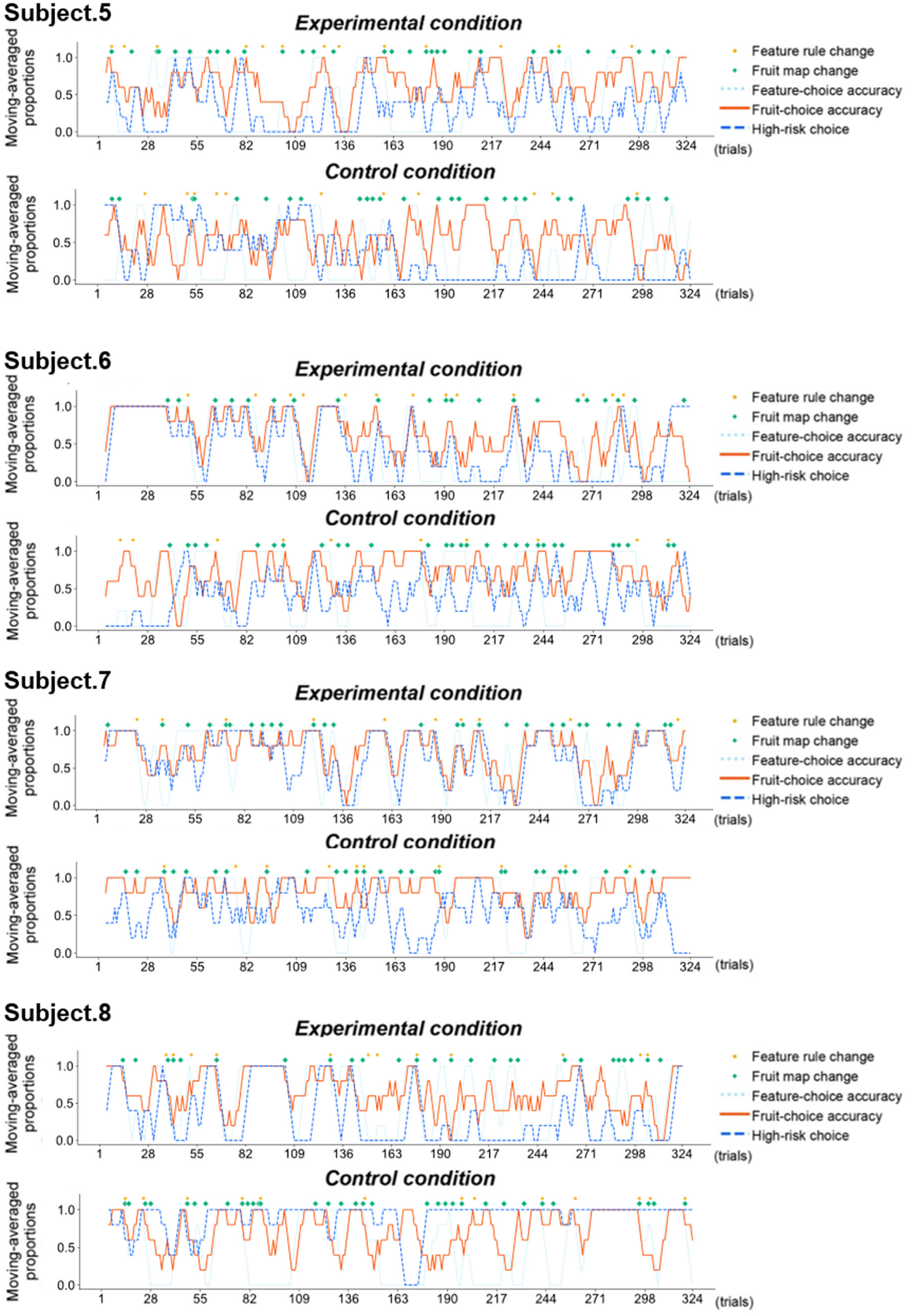
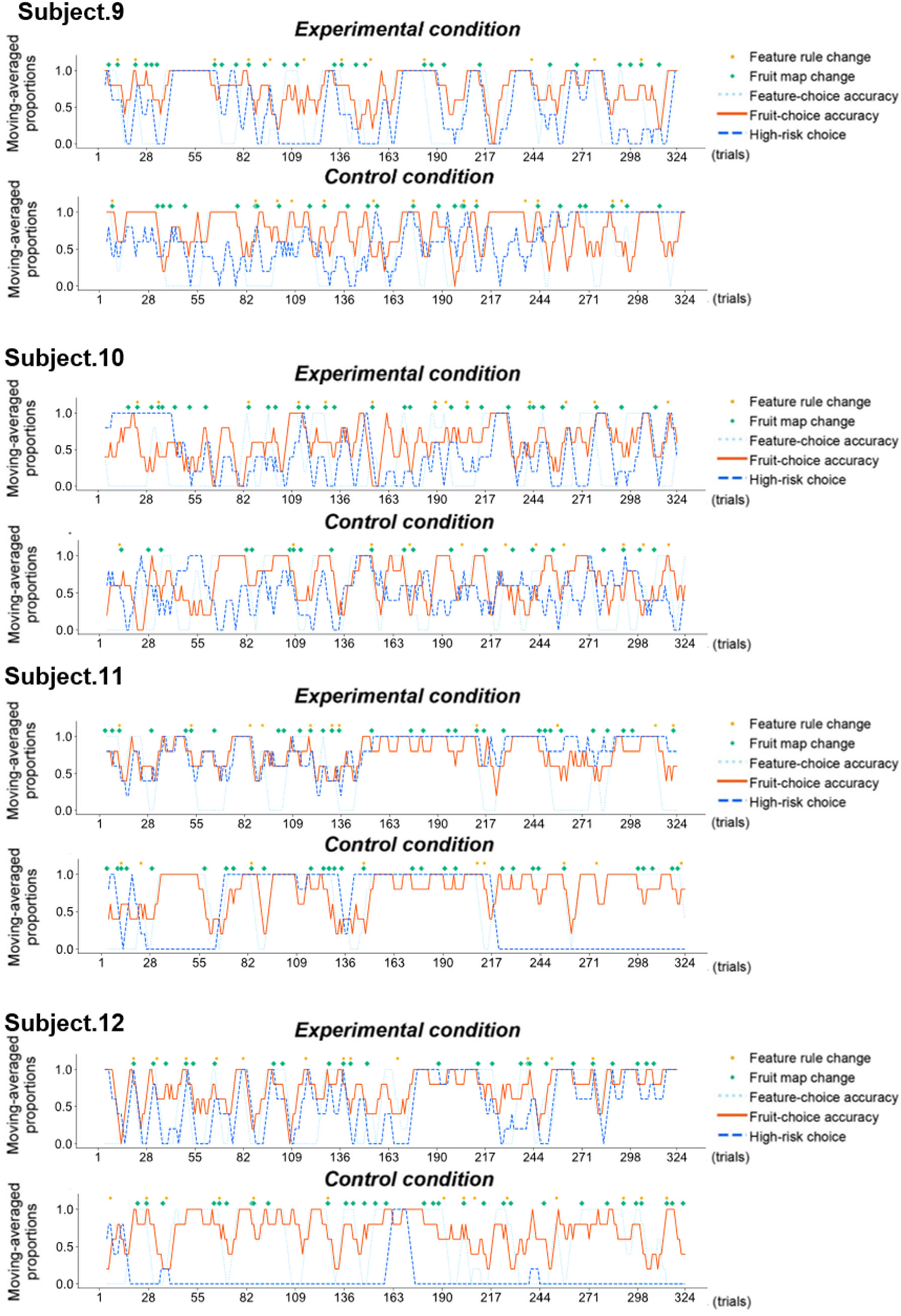
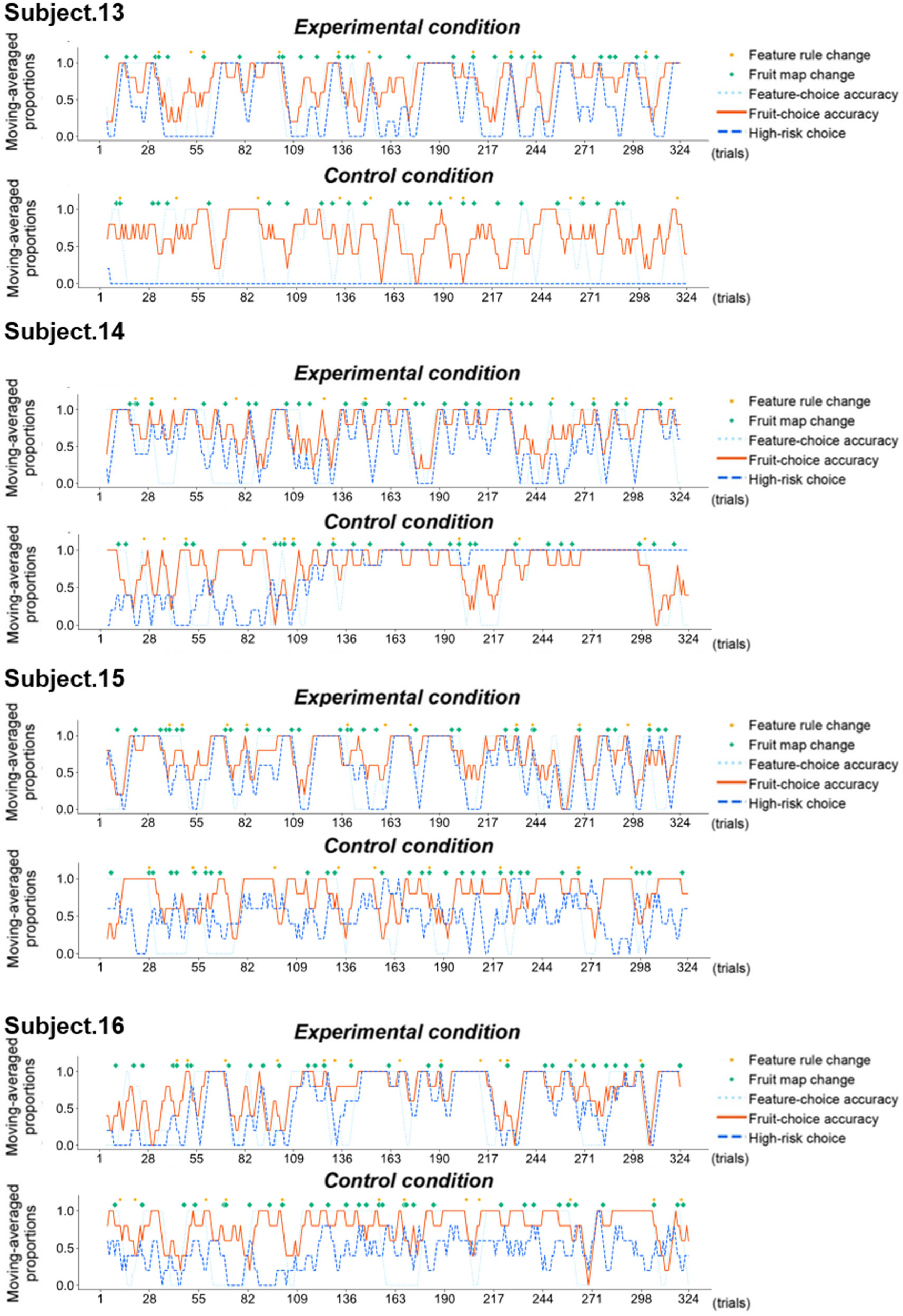
Behavioral data of each participant included in the main analysis in Experiment 1. Time courses of feature-choice accuracy, fruit-choice accuracy, and high-risk choice, smoothed using a 5-trial moving average, for each participant included in the main analysis. Correct feature choice, correct fruit choice, and high-risk choice were coded as 1 and as 0 otherwise. The upper and lower panels show the experimental and control conditions, respectively. During the task, the feature rule and the fruit mapping occasionally shifted, independently of each other, as indicated by yellow and green dots, respectively.

**Fig. S2:**
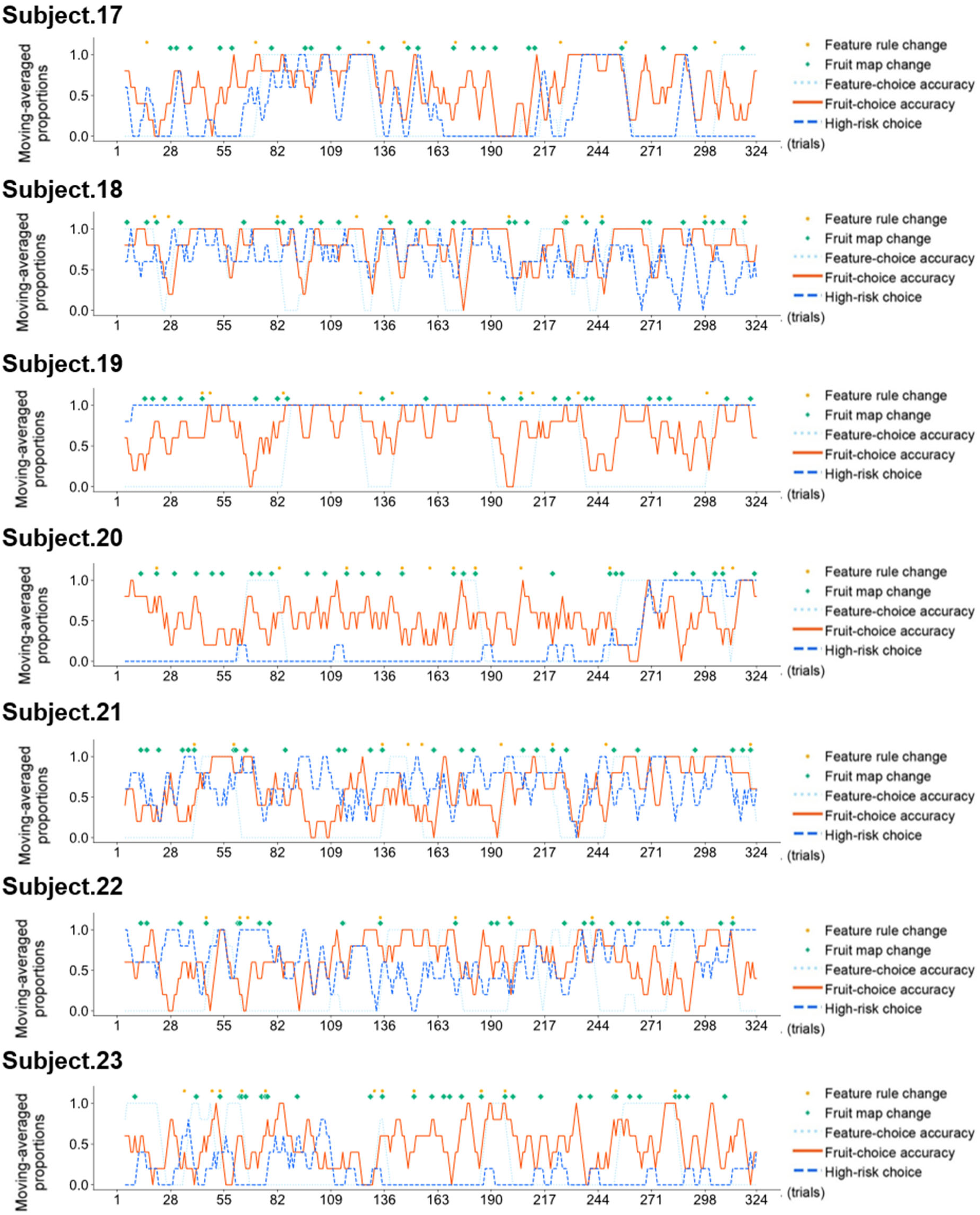
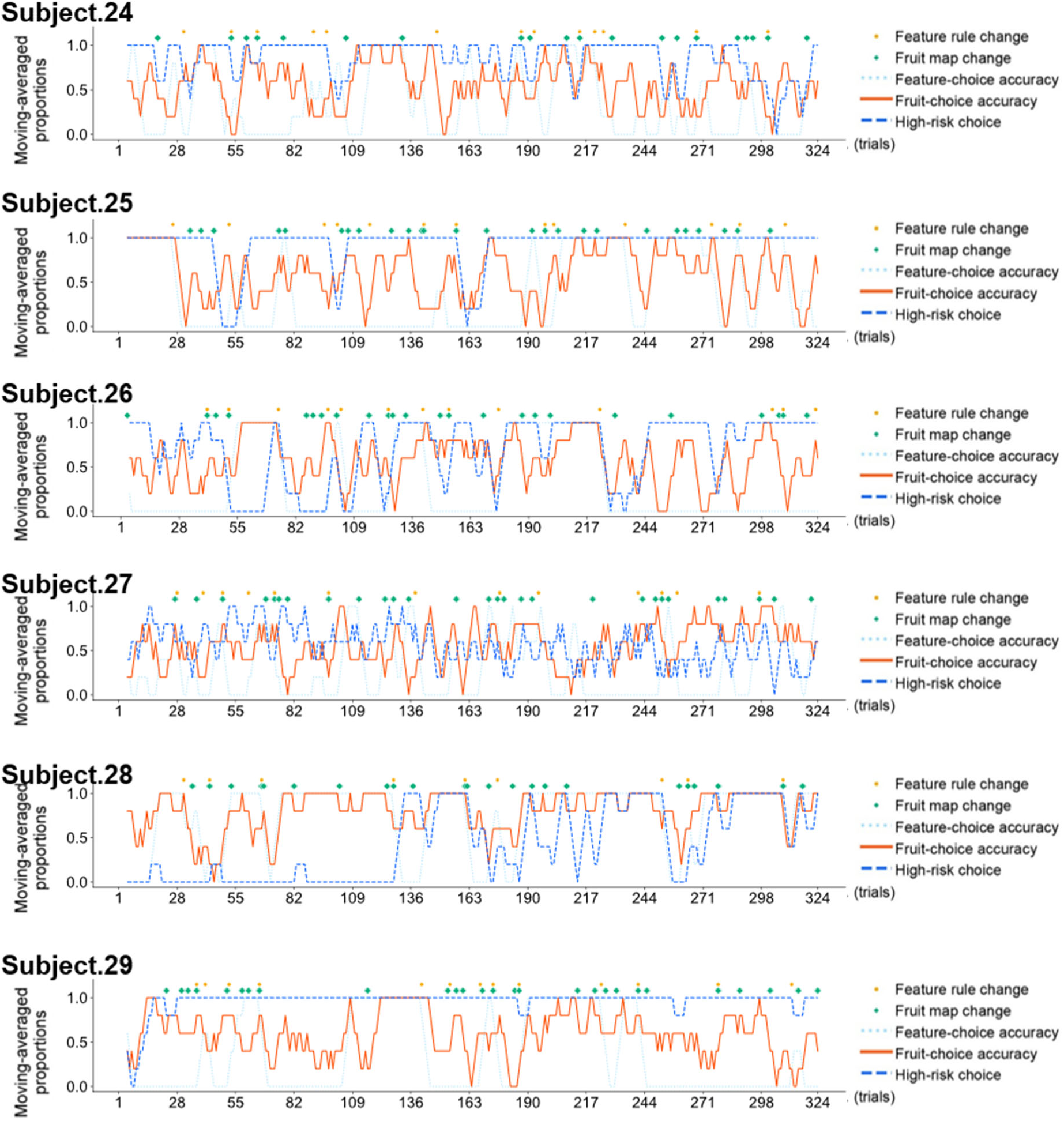
Behavioral data of each participant excluded from the main analysis in Experiment 1. Time courses of feature-choice accuracy, fruit-choice accuracy, and high-risk choice, smoothed using a 5-trial moving average, for each participant excluded from the main analysis. Correct feature choice, correct fruit choice, and high-risk choice were coded as 1 and as 0 otherwise. Only data from the experimental condition are shown. During the task, the feature rule and the fruit mapping occasionally shifted independently of each other, as indicated by yellow and green dots, respectively.

**Fig. S3:**
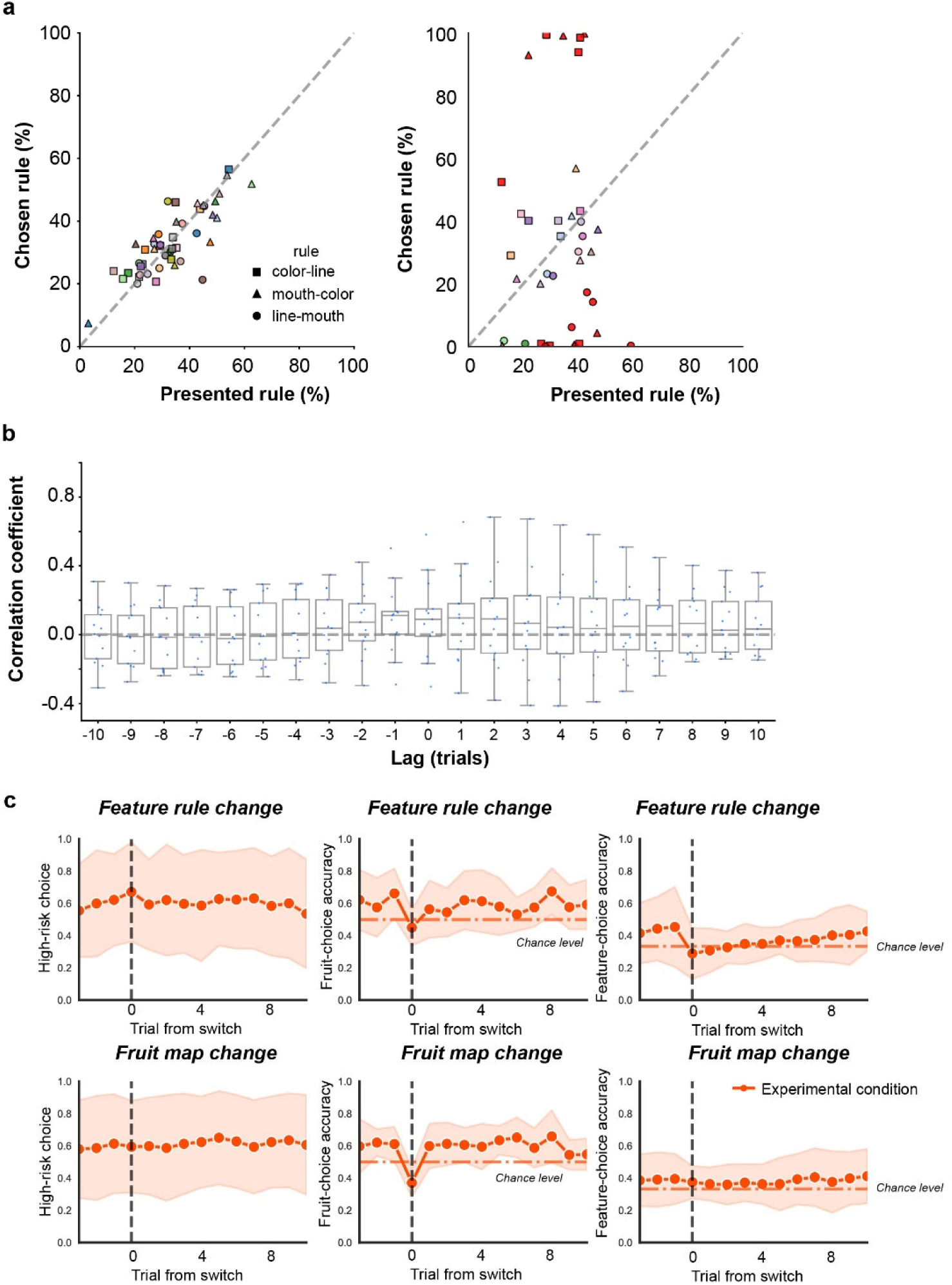
Analyses of behavioral data for excluding participants with poor task performance in the experimental condition of Experiment 1. **a,** Relations between the rate of selecting a feature rule (*y-*axis) and the rate at which the rule was correct (*y*-axis) in the experimental condition. The left and right panels are for those included in and excluded from the main analysis, respectively. Three data points are shown for each participant, corresponding to the three feature rules (color–line, mouth– color, and line–mouth). Colors on the data points vary depending on the participant except that red points indicate the participants whose rate of selecting a feature rule deviated by more than 20% from the rate at which the rule was correct. **b,** Coefficients of cross-correlation between high-risk choice and fruit-choice accuracy for the 13 participants excluded from the main analysis due to meeting either exclusion criterion. **c,** Mean time courses of high-risk choice, fruit-choice accuracy, and feature-choice accuracy around switches (shifts) in the feature rule and fruit mapping for the 13 participants excluded from the main analysis. Data were aligned to the trials at which the feature rule or fruit mapping shifted and then averaged across participants.

**Table S1.**

Participants’ answers to the post-task open-ended questionnaire in Experiment 1. Written answers were translated from Japanese into English using OpenAI’s ChatGPT (and Microsoft Copilot for comparison). The authors revised only those translations that substantially deviated from the meaning of the original Japanese writing.

### Experiment 2

**Fig. S4:**
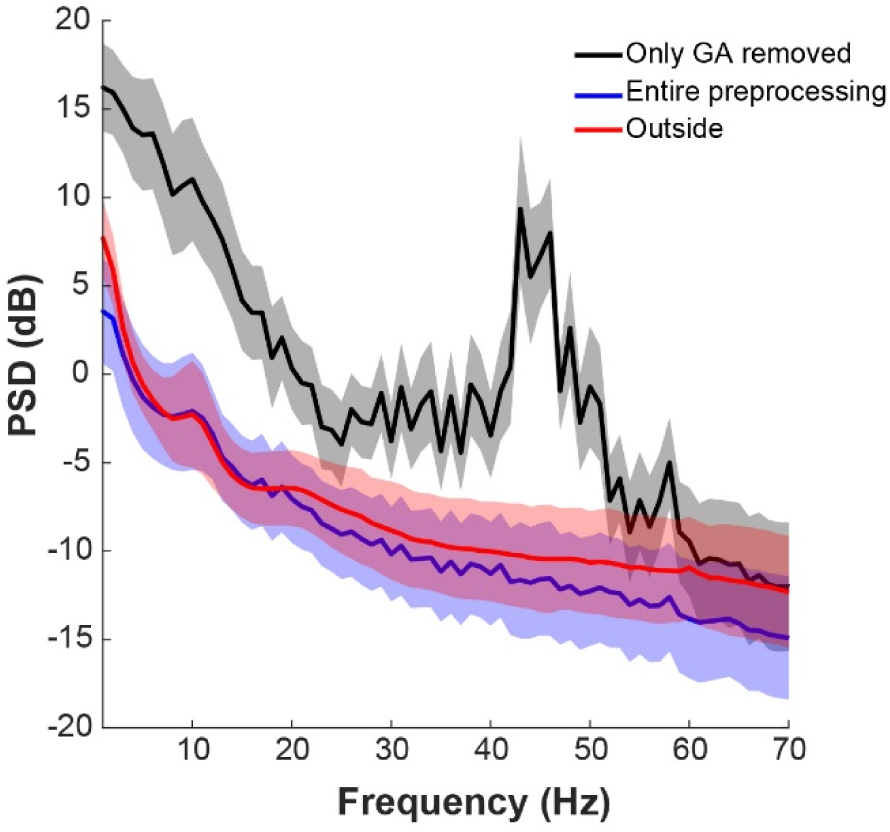
Power spectral density of EEG signals in Experiment 2. The mean power spectral density across all channels and participants, computed using EEG data. Solid lines denote the mean (black: only gradient artifact (GA) removed; blue: entire preprocessing; red: outside the MRI scanner), and the shaded regions indicate +/– 1 standard deviation across participants.

**Fig. S5:**
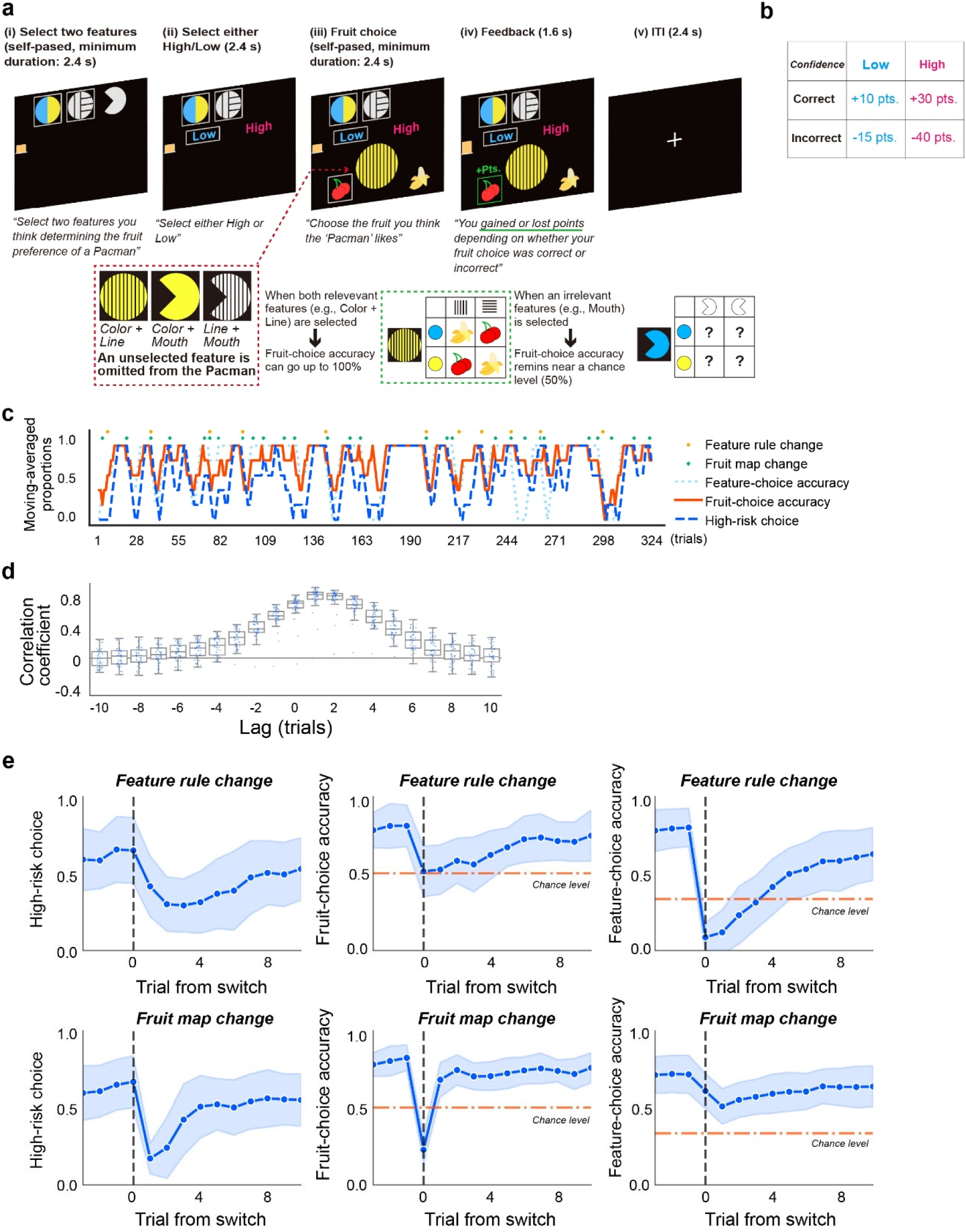
Task design and behavioral results in Experiment 2. **a**, A schematic illustration of the decision-making task conducted during simultaneous EEG-fMRI acquisition. It is similar to the one used in Experiment 1 except that the labels “High” and “Low” are added to risk options. **b**, Payoff matrix for confidence reports. High- and low-confidence choices corresponded to high- and low-risk options, respectively. **c,** Time courses of feature-choice accuracy, fruit-choice accuracy, and confidence, calculated using a 5-trial sliding window for a representative participant. The three measures exhibited similar fluctuations over the course of the task. **d**, Cross-correlation between fruit-choice accuracy and confidence. Positive correlations peaked at trial lags of +1 and +2, indicating that changes in confidence tended to follow changes in the fruit-choice accuracy. **e**, Mean time courses of confidence, fruit-choice accuracy, and feature-choice accuracy around switches (shifts) in the feature rule and fruit mapping for the 13 participants excluded from the main analysis. Data were aligned to the trials at which the feature rule (top panel) or fruit mapping (bottom panel) shifted and then averaged across participants.

### Dynamic brain state analysis using the hidden Markov model

To characterize dynamic brain states, we applied a hidden Markov model (HMM), following the procedures described in Ogawa et al. (*Sci. Rep.*, 2025). The HMM models brain activity as a sequence of discrete latent states and estimates probabilistic transitions between these states over time. Brain-state sequences were estimated from the activity of 14 predefined brain networks (Shirer et al., Cereb Cortex, 2012) derived from whole-brain fMRI data. For each network, the mean signal across all voxels within the corresponding mask was extracted. The time series were preprocessed by demeaning, regressing out six motion parameters as well as mean signals from GM, WM and CSF, and applying a temporal band-pass filter. The data were then standardized for each participant and run.

The HMM was implemented using the HMM-MAR toolbox with variational Bayesian inference assuming Gaussian-distributed observations (mean activation) and a model order of zero. The number of hidden state was set to K = 9 based on the prior study (Ogawa et al., *Sci. Rep.*, 2025). From inferred state sequences, we computed key dynamic metrics, including fractional occupancy (FO). State-specific spatial patterns were characterized by the mean activation across the 14 brain networks. To examine the relation between brain dynamics and confidence, this metric was compared between experimental conditions (e.g., high- and low-confidence).

**Fig. S6:**
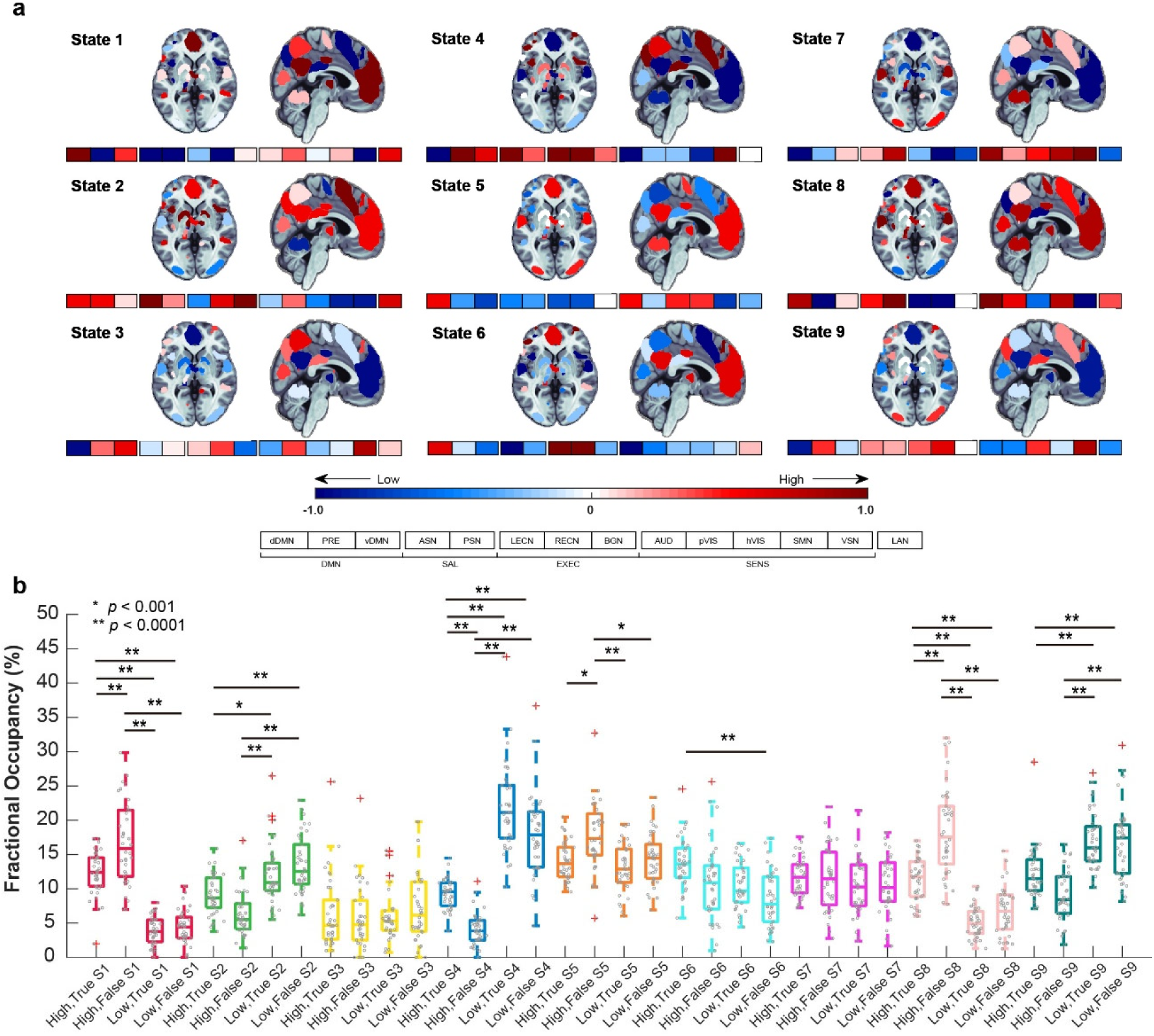
Confidence-related dynamic brain states identified in a Hidden Markov Model (HMM) analysis. **a**, Mean activation patterns of the nine hidden states identified by HMM. Each map depicts the mean activation across 14 large-scale functional networks for a given state. Network labels are shown in boxes beneath the maps. Warmer and cooler colors indicate relatively higher and lower activation levels, respectively. **b**, Fractional occupancy (FO) of each hidden state as a function of confidence and behavioral performance. FO is shown separately for high-confidence correct trials (High True), high-confidence incorrect trials (High False), low-confidence correct trials (Low True), and low-confidence incorrect trials (Low False). FO represents the proportion of the time spent in each state during the corresponding trial type. Statistical significance was assessed using one-way ANOVA followed by post hoc multiple-comparison tests with correction for multiple comparisons (*p<0.001, ** p<0.0001).

**Table S2.** Statistical results of GLM.

| **Brain regions** | **Cluster size** | **T-value** | **MNI coordinate** |  |  |
|---|---|---|---|---|---|
|  |  |  | **X** | **Y** | **Z** |
| **[Confidence] High > Low** |  |  |  |  |  |
| **R. SMA**, R. MCC, R. SFG2 | 1020 | 8.33 | 10 | 0 | 46 |
| **L. Rol. Oper.**, L. SMG, L. STG | 244 | 7.29 | -44 | -26 | 20 |
| **R. Insular**, R. Rol. Opec., R. Putamen | 260 | 6.45 | 36 | 2 | 14 |
| **R. Rol. Oper.**, R. STG, R. STG | 120 | 6.15 | 46 | -32 | 22 |
| **L. Insula**, L. Heschl, L. STG | 153 | 6.09 | 4 | -16 | -38 |
| **L. Insula**, L. Rol. Oper., L. Inf. Front. Oper. | 15 | 5.72 | -34 | 4 | 12 |
| **R. Insula**, R. Putamen, R. STG | 23 | 5.50 | 38 | -12 | 4 |
| **[Confidence] Low > High** |  |  |  |  |  |
| **R. IPL**, R. SMG, R. SPL | 2190 | 10.48 | 50 | -44 | 50 |
| **R. MFG**, R. SFG2, R. Inf. Front. Oper. | 1338 | 9.92 | 40 | 26 | 46 |
| **L. MFG2**, L. Inf. Front. Tri., L. SFG | 8 | 9.43 | -42 | 48 | 6 |
| **L. MFG2**, L. Precentral, L. Inf. Front. Oper. | 2797 | 9.35 | -46 | 16 | 44 |
| **L. Angular**, L. IPL, L. SPL | 2149 | 9.06 | -44 | -58 | 44 |
| **R. Inf. Front. Tri.**, R. Inf. Front. Oper., R. MFG2 | 40 | 6.29 | 52 | 20 | 24 |
| **R. Inf. Front. Tri.**, R. Inf. Front. Oper., R. Inf. Front. Orb2 | 13 | 6.15 | 46 | 42 | 0 |
| **R. CB Crus1**, R. CB 6, R. Fusiform | 87 | 6.09 | 46 | -70 | -28 |
| **L. SFG2**, L. MFG2, L. mSFG | 22 | 5.95 | -26 | 46 | 40 |
| **R. Caudate**, R. ACC, R. Putamen | 41 | 5.86 | 18 | 6 | 24 |
| **L. Precuneus**, R. Precuneus, R. Cuneus | 88 | 5.83 | 0 | -70 | 44 |
| **R. Precuneus**, R. SPL, L. Precuneus | 11 | 5.69 | 8 | -70 | 58 |
| **R. MTG**, R. STG, R. ITG | 18 | 5.66 | 56 | -34 | -6 |
| **R. SFG2**, R. MFG2, R. mSFG | 2 | 5.22 | 22 | 48 | 42 |

**Table S3.** Brain regions selected by sparse logistic regression.

| **Brain region** | **Cluster size** | **Peak value** | **MNI coordinate** |  |  |
|---|---|---|---|---|---|
|  |  |  | **x** | **y** | **z** |
| **R. Inf Front. Tri.** | 411 | 0.002561 | 40 | 18 | 24 |
| **L. MOG** | 133 | 0.002429 | -30 | -84 | 16 |
| **L. Fusiform** | 210 | 0.002389 | -38 | -62 | -12 |
| **R. STG** | 37 | 0.001897 | 52 | -42 | 16 |
| **L. Insula** | 51 | 0.001884 | -30 | 24 | 2 |
| **R. SOG** | 66 | 0.001847 | 22 | -70 | 34 |
| **R. Lingual** | 17 | 0.001768 | 24 | -54 | -10 |
| **R. IOG** | 110 | 0.001722 | 40 | -70 | -14 |
| **R. Fusiform** | 57 | 0.001655 | 44 | -44 | -16 |
| **R. Insula** | 75 | 0.001561 | 38 | 20 | 0 |
| **R. Fusiform** | 18 | 0.001501 | 30 | -48 | -40 |

## Appendix

Prior to the experiment, we gave participants the following instructions along with a diagram shown in Fig. 1a.

*You will perform a decision-making task. You will see a pacman-like object on a computer monitor (“pacman” hereafter). Your task is to infer the pacman’s favorite fruit from its features. The pacman consists of three features: color, line orientation, and mouth direction. Its favorite fruit is governed by a combination of two of the three features. Suppose that the combination of color and line orientation governs the pacman’s favorite fruit. For instance, a blue pacman with vertical lines may like bananas whereas a yellow pacman with vertical lines may like cherries. In this case, the third feature, namely, mouth direction, is irrelevant to the fruit preference. You need to identify the relation between the pacman’s features and its fruit preference and choose the correct fruit for a pacman image shown on the monitor*.

*Each trial runs as follows: 1) Select the two features you think are relevant to the pacman’s fruit preference; 2) choose either the light-blue or pink option; 3) choose the fruit for a pacman image shown on the monitor; and 4) see how many points you have just gained for a correct fruit choice or lost for an incorrect fruit choice. You will repeat this throughout the task. Try to find the two features governing the pacman’s fruit preference as quickly as possible to maximize the total number of points. The more points you gain, the larger the bonus payment you will get. You will make choices by pressing the up, left, and right arrow keys on a numeric keypad*.

*We now turn to details of each stage in a trial. First, you will see three icons corresponding to color, line orientation, and mouse direction. Here, you select two icons you think govern the pacman’s fruit preference using the corresponding arrow keys. When you press a key, the corresponding icon will have a white box around it. If you press the same key again, the box will disappear. Once 2.4 seconds have elapsed, you will proceed to the next stage if exactly two features are selected; otherwise, you will not proceed. Note that positions of the three icons will remain constant within a block of 54 trials but will be shuffled at a block transition. Second, select either the light-blue or pink option using the left or right key. Their points will randomly change across trials. Note that the computer will randomly select either color. Press a key to change the option if you wish. Once 2.4 seconds have elapsed, the selected color will be your choice. Third, you will see a pacman that only includes the two features you selected at the beginning of the trial. Choose the fruit you think the pacman likes using the left and right keys. Positions of the two fruits will randomly change across trials. Once 2.4 seconds have elapsed, you will proceed to the next stage when a fruit is selected. Lastly, you will see a green box if your fruit choice was correct; otherwise, a red box. Importantly, you will gain points as long as a fruit choice is correct, regardless of the selection of two features. Yet you are unlikely to make correct choices consecutively if you select a wrong feature. Suppose that color and line orientation are the correct combination of features. In this case, selection of these two features will display a pacman that helps you make a correct fruit choice. If you instead select the combination of color and mouth direction, the pacman will be missing lines. In this case, the appearance of the pacman will not help you make a correct fruit choice*.

*During the task, you will see an orange bar on the left side of the monitor. The bar indicates your current total points. You will start a block with 50 points, corresponding to the small white line at the bottom of the bar. One point is worth 1 yen. Those points will serve as a basis for the bonus money you will get in addition to base pay for participation. Specifically, the bonus will be the multiple of 50 yen closest to your total points in a block. If the total number of points is negative, the bonus will be 0 yen. The maximum bonus for today is 5,000 yen*.

*Each block will last approximately 10 minutes. You may take a short break between blocks if you wish. During a block, the correct combination of features and the pacman’s favorite fruit will each change randomly several times. When the favorite fruit changes, only the fruit preference between two fruits will be reversed while the correct combination of features will remain unchanged. A new block will be the continuation of its proceeding block. Thus, the correct combination of features and the pacman’s favorite fruit in the last trial of the proceeding block may remain the same in the first trial of the new block. Your goal is to maximize the bonus amount*.

*Before the main task, you will have three practice blocks. In the first practice block, you will see either the light-blue or pink option. Select the option displayed on the monitor. In the second practice block, the correct combination of features and the pacman’s favorite fruit will remain unchanged. Try to identify them as quickly as you can. In the third practice block, the correct combination of features and the pacman’s favorite fruit will each change once. Try to find new rules*.

